# Mechanical stress induces anatomical changes, tomato early flowering, and increased yield involving ethylene and auxins

**DOI:** 10.1101/2024.12.15.628576

**Authors:** Jenifer Castro-Estrada, Sergio M. Salazar, Julieta V. Cabello, Jorge A. Mariotti-Martínez, Raquel L. Chan, Elina Welchen

## Abstract

Plants have evolved mechanisms to perceive and face mechanical stress (MS) caused by physical forces, including compacted soils, winds, rain, pathogens, and interactions with animals and plants. Previous research indicated that applying mechanical treatment (MT) to Arabidopsis increases both xylem area and seed yield. To explore sustainable tomato production, we applied MT - combining stem bending, weighting, and particular touching - to 10-day-old seedlings, using a specific weight on the upper stem for 48 h. Two days after the treatment, we observed stem enlargement and increased the number of xylem vessels and area in MT plants. Additionally, we noticed earlier flowering, leading to increased tomato production. The transcriptome of MT-treated plants revealed significant changes in the expression of several essential genes involved in central metabolism, growth responses, and crucial phytohormone signalling. By studying different tomato mutants in the ethylene and auxin signalling pathways, we demonstrated that both hormones play essential roles in the plant responses to combined MT. Our findings suggest that combined MT generates a beneficial MS in tomato plants that induces plant morphoanatomical changes that promote early flowering and increased yield, providing a promising strategy for sustainable agriculture.

**Highlight:** A combined mechanical treatment applied to tomato seedlings enhances stem width, increases the number of vascular bundles, promotes early flowering, and improves fruit yield, involving auxin and ethylene pathways.

## Introduction

Plants are constantly influenced by the dynamic environment and exposed to different external stress factors. These factors, such as salinity, drought, and temperature fluctuations, can significantly impact their growth, development, strength, and productivity, individually or in combination, leading to substantial losses in plant productivity (Khan *et al*., 2023; Kamatchi *et al*., 2024; Ramegowda *et al*., 2024;). Aside from those mentioned above external stressing factors, along their life cycle, plants are exposed to various forms of exogenous mechanical stress (MS) that refer to the physical forces or pressures, such as dry-compacted soils, winds, rains, pathogens, animals and interactions with other plants, triggering adaptive responses that may adjust growth patterns (Mundaya Narayanan *et al*., 2024). There are also endogenous forms of MS, which play crucial roles in plant development and plant-environment interactions by influencing cellular growth and division through turgor pressure and cell wall stiffness. This stress alters the microtubule network orientation, guiding cell division and influencing plant morphogenesis (Du and Jiao, 2020; Sampathkumar, 2020). Some of these responses include the role of specific hormones interacting with each other to orchestrate a controlled equilibrium that benefits the plant’s fitness (Cabello and Chan, 2019; Khan *et al*., 2024). Such adjustments usually involve enhancement of plant resilience, improving growth, cross-stress tolerance, and nutritional value (Mundaya Narayanan *et al*., 2024).

Focusing specifically on exogenous MS factors and according to their different sources, these can include the induction of pressure gradients, gravitropic responses, and touch-induced stimuli known as thigmomorphogenesis. Plants have evolved fine-tuning mechanisms to perceive these forms of MS through specialised cells and orchestrating different responses via morphological, physiological, and biochemical adaptations (Trinh *et al*., 2021; Kouhen *et al*., 2023).

Tomatoes (*Solanum lycopersicum* L.) are among the most widely consumed and nutritious vegetables globally. They are rich in vitamin C, lycopene, beta-carotene, phenolics, organic acids, and other bioactive and health-promoting components (Quinet *et al*., 2019). Tomatoes are the second most important vegetable crop, next to potatoes, and the world produces about 100 million tons of fresh fruit from 3.7 million ha per year (FAO, 2024). According to the Tomatoes Global Market Report, the tomato market is expected to reach $239.28 billion in 2027 (ReportLinker, 2022)

Previous studies have induced mechanical stress (MS) responses in tomato plants to enhance their characteristics. Rubbing triggered a thigmomorphogenic response, leading to tissue lignification, increased antioxidant defence through up-regulation of anthocyanin synthesis genes, and reduced stem elongation (Depege *et al*., 1997; Saidi *et al*., 2009; Yoon *et al*., 2024;).

Techniques such as brushing and physical impedance also limited plant height and biomass. Additionally, wind inhibits stem growth through a thigmotropic response (Van Gaal and Erwin, 2005). These treatments strengthened stems and petioles (Garner and Bjo□rkman, 1999; Duman and Düzyaman, 2003). Controlled bending at the stem base reduced growth and height (Coutand, 2000). Overall, all these adaptations illustrate how plants respond to MS.

Stem morphology and architecture are closely related to plant fitness, resilience, and production. Previous studies have shown that wider stems and an increased number of vascular bundles are linked to high-yielding plants in Arabidopsis (Ré *et al*., 2014; Zhao *et al*., 2015; Cabello *et al*., 2016; Cabello and Chan, 2019; Colombatti *et al*., 2019). These characteristics lead to improved tolerance to flooding (Cabello *et al*., 2017; Torti *et al*., 2020) and pathogens resistance (Mencia *et al*., 2020; 2021).

Stem growth and vascular bundle multiplication are events in which several phytohormones are essential players. Previous reports involved auxin and ethylene in MS responses such as vascular cell division, cambial proliferation, secondary growth, and wood formation in several plant species (Love *et al*., 2009; Etchells *et al*., 2012; Khan *et al*., 2024). MT was previously tested in Arabidopsis and sunflower, resulting in increased stem diameter, more vascular bundles, and enhanced seed production (Cabello and Chan, 2019). To assess whether this treatment would also be effective in fruit-bearing plants, we applied it to tomato plants to determine the morphological and anatomical changes it would induce in the plants and their fruits.

In this work, we aimed to investigate the effects of a combined MT in tomato plants, focusing on the molecular mechanisms involved in developing these complex responses. For this purpose, we applied a specific weight to the upper end of 10-day-old tomato seedlings for 48 h. This treatment combined weight force, stem bending, and touching, forcing plants to return to the straight position. Transcriptome analysis indicated that the MT immediately affected the expression of crucial genes in the TOR kinase pathway, altering the balance of essential hormones involved in tomato growth and development, such as ethylene and auxins. We observed increased stem width, xylem area of vascular bundles, and earlier flowering, ultimately leading to a higher yield. Characterising mutants with altered auxin and ethylene signalling demonstrated that these hormones are critically needed for the adaptive response.

## Materials and methods

### Plant material and growth conditions

Tomato plants (*Solanum lycopersicum* (L)) from Ailsa Craig (AC), M82, and Money Maker (MM) cultivars, and the commercial cultivar Platense developed in Argentina (Amado Cattaneo, 2021), and hybrids Regina® and Chalchalero® developed in U.S.A. (BHN Research.), were used in different assays as described in the corresponding Figure legends. Tomato mutant plants in hormonal pathways were gently provided by Dr María Laura Vidoz (Plant Physiology Laboratory, FCA-UNNE, Instituto de Botánica del Nordeste, IBONE-CONICET). We evaluated mutant plants in the auxin pathway: Diageotropic (*dgt*) (auxin-insensitive phenotype) (Kelly and Bradford, 1986) and Entire (*e*) (auxin overproducer phenotype; Zhang *et al*., 2007), and in the ethylene-responsive pathway: Never ripe (*Nr*) (insensitive to ethylene; Lanahan *et al*., 1994), and Epinastic (*epi*) (ethylene overproducer; Barry *et al*., 2001).

For in-greenhouse assays, seeds were sowed on soil using 0.25 L or 0,5 L pots filled with GROWMIX® Multipro™. Experiments were carried out in parallel from 2022 to 2024 at the Institute of Agrobiotechnology, located in Santa Fe, Argentine (31°38’17.1’S, 60°40’01.8’W) and in the INTA EEA Famailla, Tucumán, Argentine (27°1’37’S, 65°38’07’W). Pots were arranged randomly at a 30 cm distance under long-day photoperiod (16/8 h light/dark cycles), with temperatures ranging from 10 °C to 40 °C. Artificial lights activated when intensity fell below 600 µmol.m[²·s[¹. Sowing dates and evaluated parameters are detailed in the Figure legends.

For the *in vitro* assays conducted on plates, tomato seeds were kept in the dark at 4 °C for nearly 5 days before being sterilised in a solution of 5 % sodium hypochlorite (NaClO) and 1 % sodium dodecyl sulphate (SDS) for 30 minutes. After washing four times with distilled water, the seeds were placed on 0.1 % agar and grown under long-day conditions (16 hours light, 8 hours dark) at 28 °C and 25 °C, with a light intensity of 400-600 µmol.m[²·s[¹ in 12 x 12 cm sterile-plastic dishes containing Murashige–Skoog medium with vitamins. Whenever special growth conditions on plates have been used (i.e., treatment with hormones or pathway inhibitors), they are specified in the figure legends.

### Application of Mechanical Treatment (MT)

Mechanical treatment (MT) was carried out on tomato plants grown on soil and *in vitro*. For soil-MT, the process involved placing a homemade device, similar to a ‘wooden clip’ weighing 1.5 or 3 g, depending on the tomato variety, on the upper end of 10-day-old seedlings with a roughly 4-5 cm stem height. The device was left on the plants for 2 days (**Supplementary Fig. S1A**). At the end of the treatment, the stem width was measured using a digital calliper (**Supplementary Fig. S1C, Fig. 1A**), and its height was measured using a ruler. These measurements were taken consecutively at different times after the treatment ended and were always analysed compared with the records taken before the treatment. Each experiment included at least 6 plants for treatment and was repeated thrice with comparable results (see **Video S1** for details). For the analysis of the impact of the MT on plants grown *in vitro* on MS 1X, 1 g % agar plates, MT was applied on 10-day-old seedlings, having a hypocotyl of 3-4 cm height, with a device of approximately 1 g, for 2 days. We incorporated a homemade device made with a 3D printer crossing the plate horizontally to prevent seedlings from detaching from the agar due to the effects of gravity plus the applied weight (**Supplementary Fig. S1B**).

**Figure 1.**
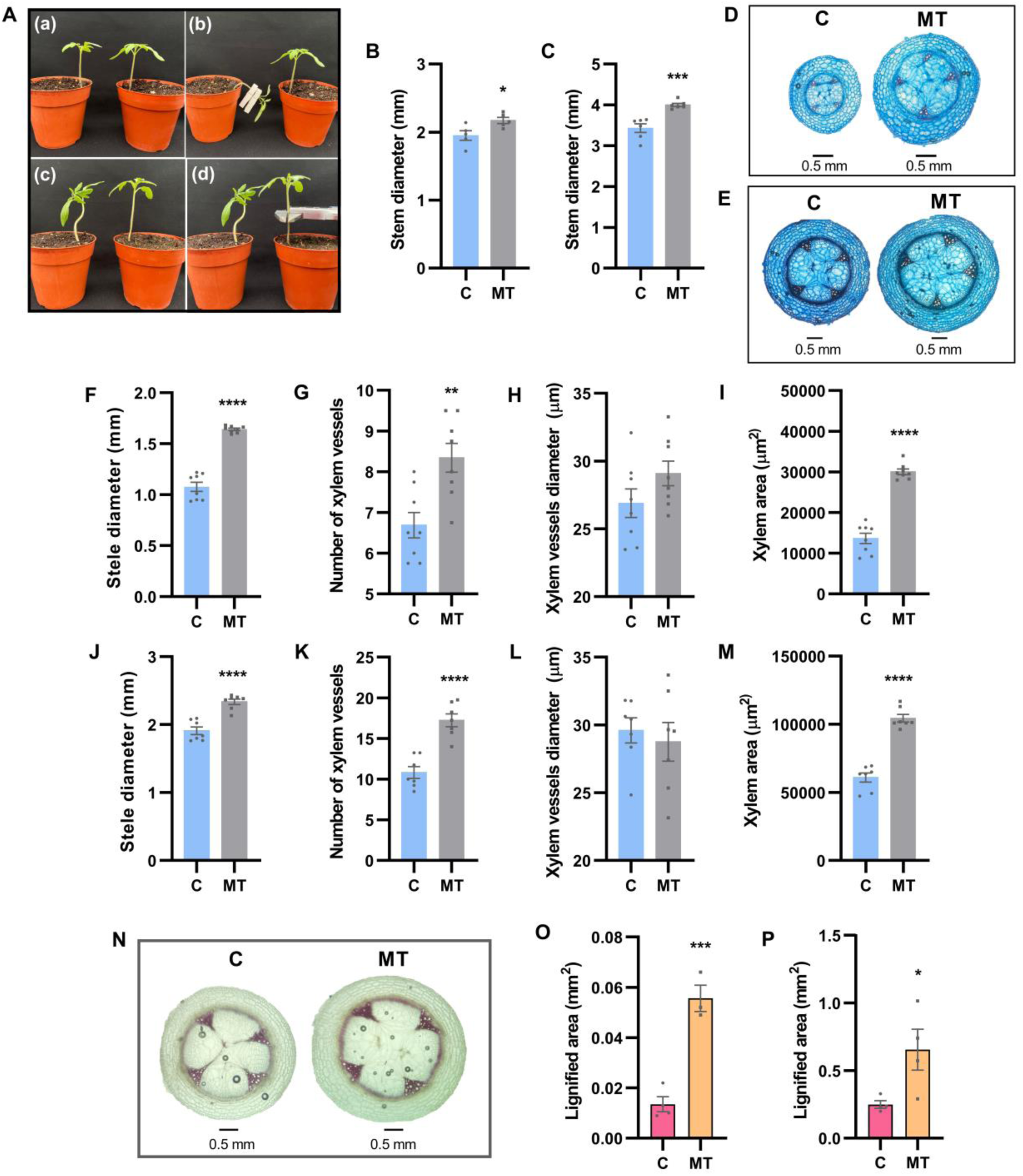
MT causes an increase in stem diameter and a modification of the vascular system in tomato plants. **(A)** Representative images illustrating the step-by-step application of MT on tomato seedlings (see video for comprehensive details): (**a**) 15-day-old seedling measuring 4-5 cm height for starting the treatment; (**b**) application of a 3 g device to the apex of the main stem for 48 hours; (**c,d**) stem diameter was measured using a digital calliper 1 cm from the base after MT application (see Video S1). The impact of MT on stem width (mm) was observed at **(B)** 48 h (48 h post-treatment (HPT) and **(C)** 10 days after treatment (10 days post-treatment (DPT). Representative images of stem cross-sections stained with astra-safranin blue after **(D)** 48 hours and **(E)** 10 days post-MT. The cross-sections were visualised using 10x lends in an optic microscope. **(F-M)** Quantitation of stele diameter (mm), number of vessels (n), xylem vessels diameter (µm), and total area of the xylem (µm^2^), performed **(F-I)** after 48 h of treatment, and **(J-M)** 10 days after treatment, for both control (C) and treated (MT) plants. **(N)** Representative images of stem cross-sections stained with phloroglucinol were shown for both control (C) and treated (MT) plants, taken 10 days post-MT. **(O-P)** Quantitative analysis of the phloroglucinol-stained sections performed **(O)** 48 h after MT began and **(P)** 10 days after the treatment ended, measuring the colour coverage by segmentation using the ImageJ software. Scale bars represent 0.5 mm. Graphs display individual values, with data presented as the mean of 10 plants ± SEM (Standard Error of the Mean). For the histological sections, data correspond to measurements from 8 randomly selected sections obtained from the total number of plants used for each experimental condition. Differences between treatments were considered significant based on an unpaired t-test at P-value (*P < 0.05; **P < 0.01; ***P < 0.001; ****P<0.0001).

### Phytohormone treatments

Tomato seedlings from different genetic backgrounds were grown for 10 days on square plates, as described above (**Supplementary Fig. S1B**). Seedlings were transferred to new plates containing MS 1x, agar 1 g % with the addition of different hormones and hormonal-pathway inhibitors as follows: 10 µM 1-aminociclopropano-1-carboxylic acid (ACC, ethylene precursor), 100 µM AgNO_3_ (Ag^+^) (ethylene-response inhibitor), 1 µM indol-3-acetic acid (IAA, synthetic auxin), or 1 µM 1-naphthyl acetic acid (NPA, auxin transport inhibitor). These different growth conditions were tested with or without applying MT to the seedlings. After 48 h of growing under these tested conditions, the tomato stem width was measured with a digital calliper (**Supplementary Fig. S1C**).

### Plant phenotyping

Different plant traits were scored on control (C) and weight-treated (MT) tomato plants, including stem height and width, flowering time, fruit number, fruit quality, and plant yield. Measurements were performed manually with a ruler or digital calliper. All the experiments were conducted with 16 plants per treatment and repeated at least four times. Stented stem histological cuts were photographed for a detailed analysis of stem anatomy. We used the ImageJ free software (Schneider *et al*., 2012) to evaluate parameters such as xylem area, stele diameter, vascular bundle and xylem vessel number and size.

### Analysis of fruit quality/Fruit quality measurements

The freshly harvested fruits were used to evaluate the impact of the MT on fruit quality. Total soluble solids were terminated at the harvest time from the tomato juice, using a refractometer (ATAGO, PAL-BX ACID2) and recording three readings per fruit on randomly chosen 5 fruits from 6 plants (Furio *et al*., 2022). Acidity was determined using an aliquot of 10 g of tomato juice in 100 mL of deionised water. It was determined using a refractometer (ATAGO, PAL-BX ACID2). Titratable acidity was expressed in g of citric acid per L (Sapers, 1994). The surface colour was evaluated with a colourimeter (Minolta, Model CR-300, Osaka, Japan) by measuring the parameters L*, a* and b*. Negative L* indicates darkness, and positive L* indicates lightness. Negative a* indicates the green colour, and positive a* indicates the red colour. A high positive b* indicates a more yellow colour, and a negative b* indicates a blue colour. The chroma value (C*), calculated as C* = (a*2 + b*2)1/2, indicates the intensity or saturation of the colour. The colour was measured in three random positions for each fruit (Agüero *et al*., 2015). Thirty fruits were analysed for each treatment, and the experiments were conducted in triplicate.

### Histological analysis of stem-cross section

Tomato stems were fixed in 70 % ethanol for 48 h. Freehand cross-sections were prepared under a stereo microscope (Leica EZ4) on slides (25.4 x 76.2 mm) using commercial razor blades. The sections were rinsed with 50 % bleach, followed by five washes with distilled water, then stained using a successive double staining method with astra-safranin Blue (Arend *et al*., 2008). After staining, the sections were washed with distilled water to remove any excess dye and then mounted in a 1:1 mixture of water and glycerin (Rodrigues Marques and Kasue Misaki Soares, 2022). Microscopic optic visualisation (Eclipse E200, Carl Zeiss, Axiostar Plus, Göttingen, Germany) using 10x lends was conducted to determine the following variables: stem, stele, pith, and cortex diameter (µm), number of xylem vessels, and xylem area (µm^2^). Images were analysed using the ImageJ (Schneider *et al*., 2012) program, employing freehand tools for the xylem area and line selection for stele and xylem vessel diameter measurements.

### Lignin detection and quantitation

To detect and quantify lignin deposition, cross-sections of plant stems subjected to MT and their respective controls were made using a sharp blade with the freehand technique. Afterwards, they were stained with 1 % phloroglucinol, following a method described by (Rodrigues Marques and Kasue Misaki Soares, 2022). The stained cell walls of lignified sections were flamed, and 25 % hydrochloric acid was added to give them a purplish-red colour. Finally, the images were analysed using the free ImageJ software (Schneider *et al*., 2012). A colour segmentation coverage measurement was performed using the same threshold for all cases.

### Determination of starch and soluble sugars

Determination of starch and soluble carbohydrates was performed according to Canal *et al*. (2024). Briefly, 30 mg of leaves and 50 mg of stem pulverised tissues were resuspended in 250 μl of 80 % ethanol in 10 mM Hepes-KOH (pH 7.0) and incubated for 20 min at 80 °C. Subsequently, the mixture was centrifuged at 13000 rpm for 5 min. The supernatant (S1) was carefully transferred to a new tube, while the pellet was resuspended in 250 μl of 50 % ethanol in 10 mM Hepes-KOH (pH[7.0), incubated at 80 °C and centrifuged at 13[000[rpm for 5[min. Next, the pellet was resuspended in 150 μl of 80 % ethanol in 10 mM Hepes-KOH (pH[7.0), followed by incubation at 80 °C and centrifugation at 13000 rpm for 5 min. The resulting supernatant (S3) was added to the combined S1 and S2 supernatants, and an aliquot was used to determine soluble carbohydrates (Stitt *et al*., 1989). The pellet saved in the ethanolic fraction was used to quantify starch content using an enzymatic method (Hendriks *et al*., 2003).

### Determination of total chlorophyll content

The chlorophylls were determined by measuring the absorbance at 645 and 665 nm in a microplate with 96 wells containing 50 μl of the ethanolic extract obtained as the final step of the previous protocol (S1+S2+S3) diluted with 120 μl of absolute ethanol. The following formula was used to calculate the amounts of Chl*a* (Chl(μg/well) = 5.21*A_665_–2.16*A_645_) and Chl*b* (Chl (μ/well) = 9.29*A_645_–2,74*A_665_).

### RNA isolation and analysis

RNA was isolated as described by Welchen *et al*. (2012). Relative transcript levels were measured by reverse transcription and quantitative PCR (RT-qPCR) using specific primers. Reverse transcription was performed on 1 μg total RNA using an oligo (dT)_18_ primer and MMLV reverse transcriptase (Promega). For qPCR, an aliquot of the cDNA was subjected to amplification with specific primers (**Supplementary Table S1**). Amplification was detected in a Bio-Rad CFX96 thermocycler using SYBR Green. Quantifying mRNA levels was achieved by normalising the transcript levels against β*-2-TUBULIN* (Solyc10g086760; Albuquerque *et al*., 2021) following the ΔΔCt method. All reactions were performed with at least three biological replicates, and the bars represent the mean ± SEM. All primers used for RT-qPCR analyses are listed in **Supplementary Table S1**.

### RNA-seq analysis

For RNA-seq analysis, total RNA was extracted from the stem of control (C) and treated (MT) 10-day-old seedlings grown on soil. Three biological replicates from each genotype were used for RNA sequencing using an external service (BGI Americas Corporation, Cambridge, MA, USA, Project number: F23A480000788-01_SOLaswfR) and DNBSEQ^TM^ technology with a 100-bp read length. Raw data were filtered with SOAPnuke v1.5.2 (Chen *et al*., 2018), and clean reads were cleaned and aligned to the *Solanum lycopersicum* ITAG4.0 reference genome (solgenomics.net) using HISAT2 v2.0.4 and Bowtie2 v2.2.5. Analysis was conducted using Dr Tom’s web-based solution (https://www.bgi.com/global/service/dr-tom). Differential gene expression of MT versus C samples was evaluated using DEseq2 (Love *et al*., 2014). The phyper function in R software is used for enrichment analysis, estimating the *P*-value, and the *Q*-value was obtained by correcting the *P*-value. Generally, *Q*< 0.05 was considered as the significant enrichment.

### Western blot analysis

Western blot analysis was performed according to Canal *et al*. (2024). Proteins were denatured at 95 °C for 5 min and then centrifuged at 12000 g for 10 min. Subsequently, 10 μl of the supernatant was separated on a 10 % SDS-PAGE and transferred to a polyvinylidene difluoride membrane. Membranes were hybridised with rabbit antibodies against P-RPS7 (AS 194302, Agrisera; dilution 1:10000), RPS7A (AS194292, Agrisera; dilution 1:10000), or actin (AS 132640, Agrisera; dilution 1:10000), followed by detection with anti-rabbit IgG secondary antibodies conjugated with horseradish peroxidase (HRP) (Agrisera; dilution 1:50000) and SuperSignal West Pico Chemiluminescent Substrate (Thermo Fisher Scientific). Rat anti-HA antibody (11867423001; Roche; dilution 1:10000) was detected using HRP-conjugated anti-rat IgG (Agrisera; dilution 1:20000).

### Statistical analysis

The statistical analysis used is specified in the legend of each corresponding figure. Unless indicated in a particular study, data were evaluated by single-factor ANOVA (with Tukey’s or Holm-Sidak’s post-hoc test, P < 0.05). Student’s unpaired t-test was used to compare the two groups, considering a significance value of P < 0.05.

## Results

### MT increases stem diameter and xylem area in tomato plants

Aiming to understand how MS affects the development of tomatoes, we carefully designed a combined treatment with a simple “clip-type” device placed on the top of the plant stem (**Supplementary Fig. S1A, Video S1**). Such specific application generated a MS that included multiple responses against the combined effect of touching the stem, bending it with weight, and subjecting it to mimicking a kind of “microgravity”. Firstly, given their anatomical differences, we assessed the required weight and developmental stage to provoke the MS responses for each tomato variety. For Ailsa Craig (AC) tomatoes, we found that the most significant effect of the MT was achieved by using a 3 g device for 48 h on the apex of the main stem of 10-day-old seedlings exhibiting 4-5 cm-height ( **Fig. 1A (a, b)**). After 48 h from the start of the MT treatment (48 HPT, hours post-treatment), plants responded by displaying opposite forces to return to their natural upright position (**Fig. 1A (c)**). Immediately after it ended, the stem width was measured 1 cm from the base using an electronic calliper (**Fig. 1A (d)**). Treated plants (MT) significantly increased their stem width by around **15%** compared to untreated plants used as controls (C) at 48 HPT (**Fig. 1B**), and the impact of the MT on the stem width persisted at least 10 days after the treatment ended (10 DPT, days post-treatment) (**Fig. 1C**).

To assess morphological changes in depth conducting to stem enlargement, stained stem cross-sections from MT plants and untreated controls (C) were microscopically analysed at 3, 6, 12, 24, and 48 h post-treatment initiation (**Supplementary Fig. S2**). The visualisation revealed that changes in stem structure began at 6 h (**Fig. 1D, Supplementary Fig. S2**) and persisted almost 10 days post-MT (**Fig. 1E**). Stele diameter and the number of xylem vessels significantly increased, leading to a substantial increment in the stem xylem area at 48 h from the MT initiation (**Fig. 1F-I**). Remarkably, these significant increases persisted almost at 10 DPT (**Fig. 1J-M**). Furthermore, analysing cross-sections of MT stained with phloroglucinol (**Fig. 1N**) showed that stems of the MT-exposed plants underwent a notable lignification after 48 h of treatment initiation, and these increases persisted at least 10 days after it ended (**Fig. 1O, P**).

The observed results indicated that the plant responses to the MS imposed became noticeable 6 h after its initiation. Such impact was characterised by the enlargement of stem diameter, the area of the conducting vessels, and the lignin deposition on stem cell walls. Although the weight and treatment period should be adjusted for each tomato variety, similar results were observed on tomato varieties such as M82, Money Maker, and the Argentinian-market-available hybrid varieties Chalchalero®, Platense and Regina® (**Supplementary Fig. S3**). For example, for the commercial hybrids, we needed variable weight devices between 1.5 g to 3 g, depending on the experiment and the growth condition, and the same 48 h to achieve the maximal effect (**Supplementary Fig. S3**). These additional results in commercial hybrids indicated the broad applicability of our research.

### The MT affects the progression of the life cycle, leading to early flowering and increased fruit yield

To further investigate the impact of stem enlargement and vascular bundle multiplication, we monitored the characteristics of tomato MT plants throughout their entire life cycle. Thus, AC tomato plants grown in a greenhouse showed an increase in the average number of inflorescences between seven to eleven weeks after sowing (WAS) when subjected to MT (**Fig. 2A**). Treated (MT) plants had 100% more fruit buds by week eleven than untreated or Control (C) plants (**Fig. 2B**). The results showed that MT significantly affected the growth and development of tomato plants, increasing the number of fruit (**Fig. 2C**). Furthermore, the weight of harvested tomatoes between weeks sixteen and eighteen from sowing was around 80 % more in treated than in untreated plants (**Fig. 2D**). Although the differences between MT and controls diminished after the first flowering, the whole yield increase was approximately a 22 % higher considering total fruit number (**Fig. 2E**), and a 15 % taking into account fruit weight (**Fig. 2F**). Remarkably, the positive impact of MT on accelerating flowering and increasing production did not compromise the phenotype and quality of the fruits. The fruit number, but not the visual appearance (see **Fig. 2G** as a representative), shape, size, and fruit colour (**Fig. 2H-J**) were comparable between MT and C plants. The quality indicated by sugar content (°Brix) (**Fig. 2K**) was also similar. Moreover, the fruit “acidity”, a parameter especially desired for industrial applications because it enhances fruit preservation, guaranteeing a desirable organoleptic profile, was slightly increased in MT-treated plants (**Fig. 2L**). These findings highlight the potential benefits of early intervention with MT for maximising crop productivity.

**Figure 2.**
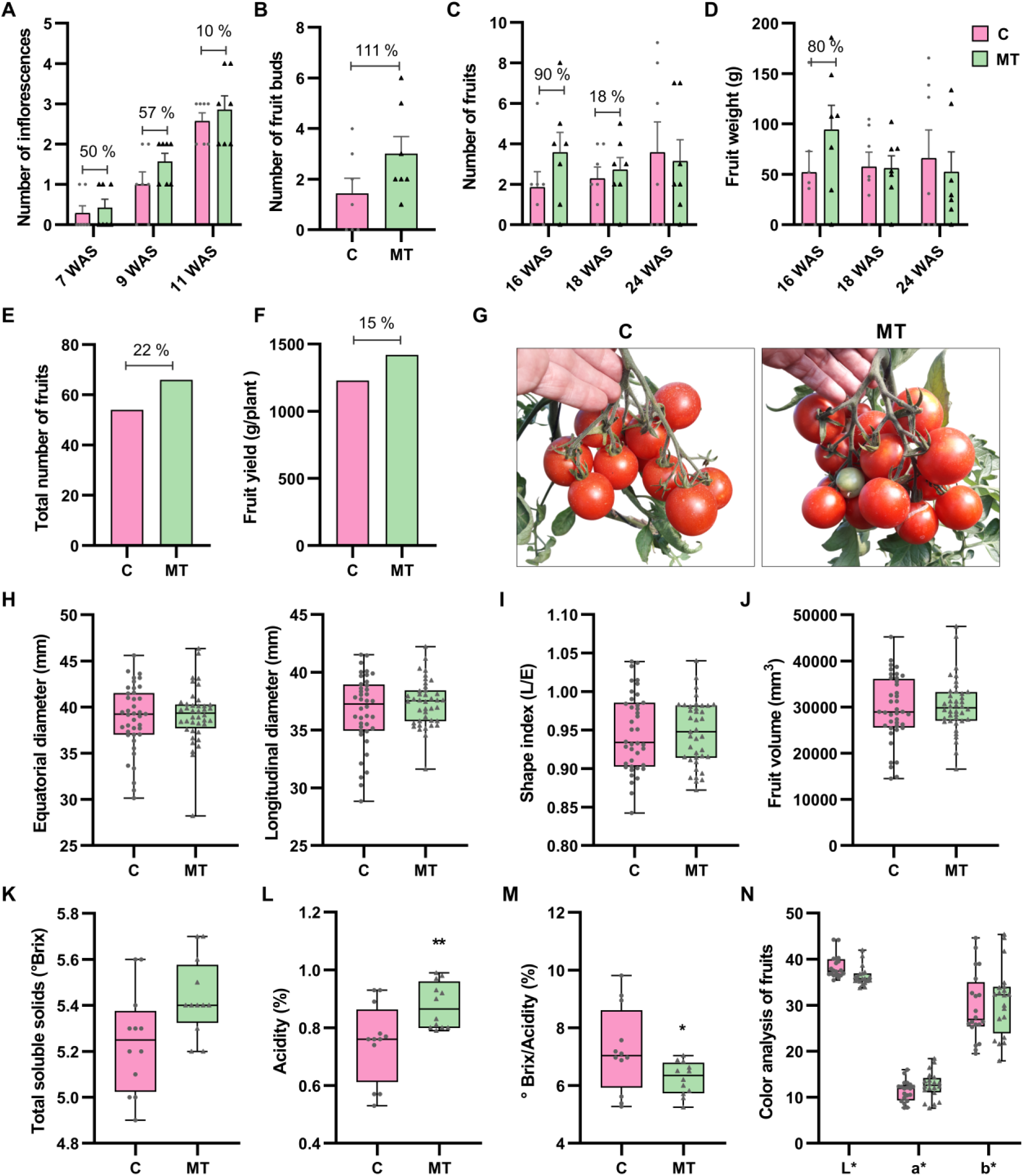
MT accelerates flowering and enhances tomato production. **(A)** The number of inflorescences in plants aged 7 to 11 WAS (Weeks After Sowing), comparing control (C) and treated plants (MT), is represented in pink and light green, respectively. **(B)** Number of fruit buds in 11-week-old plants in the C and MT groups. **(C)** Comparison of the number (n) of fruits harvested and **(D)** the weight of fruits (gr) collected from 16 to 24-week-old plants (WAS) in the MT group (light green) compared to the control group (C) (pink). **(E)** Total number of fruits harvested at the end of the life cycle in C and MT plants. **(F)** Comparison of the total fruit yield (gr/plant) obtained in C and MT plants by the end of the life cycle. **(G)** A comparative image of fruits harvested during the first weeks illustrates the differences in the number of fruits between MT and C plants. **(H-J)** Physical measurements of fruit quality, including equatorial and longitudinal diameter (mm) **(H)**, shape index **(I)**, and average volume (mm^3^) **(J)**. **(K-M)** Physiological parameters related to fruit quality: total soluble solids (°Brix) **(K)**, titratable acidity (%) **(L)**, and °Brix/acidity ratio **(M)**. **(N)** Colour analysis based on L* (lightness), a* (green-red), and b* (blue-yellow) visual parameters. The graphs display individual values, with means representing the average of 7 plants ± SEM. Differences between treatments are considered significant if determined by a t-test with a P-value (*P < 0.05; **P < 0.01).

We made similar observations with the commercial hybrid variety, Chalchalero®. We found that the stem diameter increased significantly 48 h after the treatment began (**Supplementary Fig. S4A**). As we observed with the AC variety, Chalchalero® tomatoes also exhibited an increased number and biomass of fruits (g) per plant, with significant growth noted between weeks 28 and 38 WAS (**Figure S4 B, C, F**). These weekly results were translated into an overall increase in the total number of fruits (**Figure S4D**) and the overall tomato yield per plant (**Figure S4E**) by the end of the productive period. These results were consistently reproduced, with minor differences between the years, across three growing seasons in tomato plants cultivated in greenhouses and experimental fields in San Miguel de Tucumán, Argentine.

### MT induces changes in gene expression and alters plant metabolism to increase fruit yield

We performed a comparative RNA-seq analysis on AC stems subjected to MT to unravel the molecular mechanisms involved in the significant anatomical changes observed. Samples were collected 6 h after the MT started (MT_6h) and ten days after treatment (MT_10D), together with their respective control samples from untreated seedlings (**Fig. 3A**). The analysis revealed significant changes in the expression of 2019 genes at 6 h (*Q* < 0.05); 1229 were upregulated (log_2_FC > 0.6), and 790 were downregulated (log_2_FC < −1). At ten days after MT, 412 genes showed differential expression (*Q* < 0.05), including 275 upregulated (log_2_FC > 0.6) and 137 downregulated (log_2_FC < −1) genes (**Fig. 3B**). This large number of modulated genes correlated with the significant changes observed in the phenotype of MT plants.

**Figure 3.**
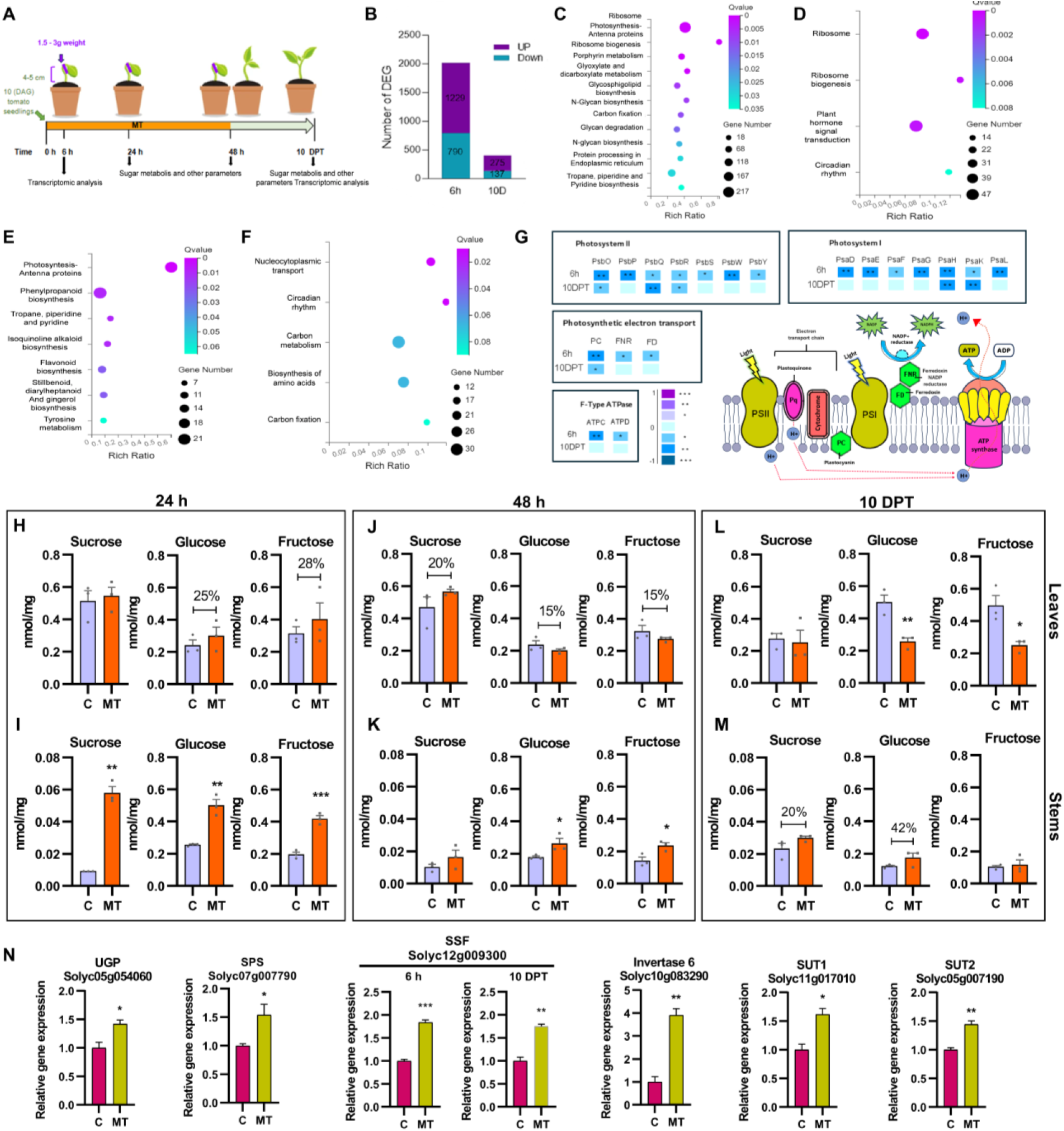
MT induces changes in gene expression and alters sugar metabolism. **(A)** Schematic representation of the analysis performed on C- and MT- groups of plants. For RNA-seq analysis, stem sections were collected at 6 h after the start of the MT (6 h) and 10 days post-treatment (10 DPT). For sugar metabolism analysis, stems and leaves in the same groups of plants were collected at 24 h, 48 h, and 10 DPT. **(B)** Number of genes statically differentially expressed (DEGs) (Q value < 0.05) after the RNA-seq analysis at 6 h and 10 DPT. Upregulated genes are shown in purple, while downregulated ones are in magenta. **(C-F)** Gene ontology (GO) enrichment analysis of the DEGs was performed according to the KEGG pathways. The analysis was conducted using Dr Tom’s web-based solution (https://www.bgi.com/global/service/dr-tom). The “y” axis displays statically enriched pathways based on the GO annotation classification, while the “x” axis represents the Rich ratio (ratio of enriched DEGs to the background genes). The colour of the circles represents the statistical significance, and the size of the circles refers to the number of genes that are overrepresented in each pathway. Qvalue <= 0.05 are considered as the significant enrichment. **(C)** GO analysis of the DEGs at 6 h after the initiation of treatment. GO pathways enrichment analysis of differentially **(D)** upregulated genes (log_2_FC>0.6) and **(E)** downregulated (log_2_FC<−1) genes at 6 h. **(F)** GO analysis of the pathway enrichment for DEGs at 10 DPT. **(G)** Schematic representation of the impact of the MT on the photosynthetic pathway and ATP synthesis: Components of photosystems I and II (PSI and PSII) and functional proteins essential for the photosynthetic machinery, photosynthesis light reactions, and ATP-synthase. Gene expression was symbolised as coloured boxes representing the log_2_ fold change of upregulated (purple) or downregulated (blue) genes. Statistical analysis of the gene expression was conducted using DESeq2 (Dr. Tom’s web-based tools). Statically significant differences were indicated with an asterisk (P < 0.05). Within each colour category, the intensity varies according to the P value (*P < 0.05; **P < 0.01; ***P < 0.001), with darker colours indicating higher levels of statistical significance. The figure was adapted from the KEGG pathway (Okuda *et al*., 2008). See **Supplementary Fig. S7** for individual graphs of each gene’s graphs generated from the FPKM (Fragments Per Kilobase Million). **(H-M)** Quantification (nmol/mg) of glucose, fructose and sucrose in the leaves **(H, J, L)** and stems **(I, K, M)** of treated plants (MT, orange) and controls (C, violet) was conducted at 24 h, 48 and 10 DPT. **(N)** Relative expression levels of induced genes after 6 h of treatment (MT), coding for genes involved in the synthesis of cytosolic glucose and sucrose (*UGP, SPS, SSF, INVERTASE*) and sucrose transporters (*SUT*). Gene expression in control plants is indicated in purple, while expression in MT plants is symbolised in light green. The graphs show individual values, with data presented as the mean of 3 plants ± SEM. Differences between treatments were considered significant according to an unpaired t-test at P < 0.05. Abbreviations: UGP: UTP GLUCOSE 1 PHOSPHATE URIDYLYLTRANSFERASE, SPS: SUCROSE PHOSPHATE SYNTHASE, SUS: SUCROSE SYNTHASE, FBP: FRUCTOSE-1,6-BISPHOSPHATASE CYTOSOLIC, SUT: SUCROSE TRANSPORTER.

We conducted a gene ontology (GO) enrichment analysis of the differentially expressed genes (DEGs) to identify the biological pathways affected by MT. Thus, at 6 h, the significant categories (*Q* < 0.05) included genes associated with ribosome biogenesis and function, biosynthesis of the photosynthetic machinery, and the synthesis and processing of secretory and membrane proteins. Additionally, there were changes affecting carbon fixation and glyoxylate metabolism (**Fig. 3C**). The analysis indicated that the most significantly upregulated pathways were associated with ribosome biogenesis, plant hormonal signal transduction, and circadian clock regulation (**Fig. 3D**). Conversely, pathways involved in photosynthesis and the biosynthesis of secondary metabolites and phenylpropanoids were found to be the most strongly downregulated (log_2_FC<-1) (**Fig. 3E**). Ten days after the treatment, when morphoanatomical changes were still observed (**Fig 1J-M**), DEGs were found in many shared categories, which again included circadian cycle, photosynthesis, and carbon metabolism genes (**Fig. 3F**).

Based on the morphoanatomical responses observed, we focused on the processes involved in generating and transforming energy available for growth. Surprisingly, MT suppressed chlorophyll biosynthesis genes after 6 h, with several changes persisting even 10 days after MT (**Supplementary Fig. S5**). Notably, such repression did not impact chlorophyll content (**Supplementary Fig. S6A**). In the same way, MT caused the repression of genes encoding components of Photosystems I and II (PSI and PSII), those relevant for electron transport and chloroplast functional photosynthetic machinery proteins such as PLASTOCIANIN (PC), FERREDOXIN-NADP REDUCTASE (FNR), and F-type ATPase (**Fig. 3G** and **Supplementary Fig. S7C, D**). Additionally, the genes encoding the LIGHT-HARVESTING COMPLEXES (LHCs) components were downregulated after 6 h of MT, persisting altered even after 10 days.

Connected to this process, we also observed reduced expression of genes involved in carbon fixation and reduction steps of the Calvin cycle (**Supplementary Fig. S8**), including those coding chloroplast RUBISCO (RBCS), RUBISCO Activase (RA), Phosphoglycerate Kinase 1 (PGK-1), and Glyceraldehyde Phosphate Dehydrogenase (GAPDH) (**Supplementary Fig. S8B, C**). Additionally, we found that other genes coding for enzymes involved in sucrose, glucose, and fructose production in the cytosol, like Fructose-1,6-Bisphosphatase (FBP) and FBA were also downregulated (**Supplementary Fig. S8E**).

To assess whether the observed gene modulation effectively affected carbohydrate content, we quantified glucose, fructose, and sucrose in stems and leaves of MT and control plants. While the soluble sugar content in the leaves of MT plants showed slight changes compared to the controls at 24 and 48 h after treatment began (**Fig. 3H, J**), the stems of the same treated plants exhibited a significant increase in glucose, fructose, and sucrose levels (**Fig. 3I, K**). This increment could indicate a response for transporting the needed sugars from the stem to the leaves to sustain the growth-promoting action elicited by the MT. However, the incremented growth in the stem width caused a reduction in the soluble sugar content (glucose and fructose) in the leaves 10 DPT (**Fig. 3L**), while starch levels did not change significantly due to MT (**Supplementary Fig. S6B**). In this sense, the expression of cytosolic glucose and sucrose synthesis encoding genes *UTP-GLUCOSE-1-PHOSPHATE URIDYLYLTRANSFERASE (UGP), SUCROSE SYNTHASE* (*SUS*) and *SUCROSE PHOSPHATE SYNTHASE (SPS)*, *INVERTASE* (*INV*), and those encoding for *SUCROSE TRANSPORTERS* (*SUT*) were induced 6 h after the treatment started (**Fig. 3N**), which could indicate the active transport process of sugars from stems to leaves.

Additional data supporting the decrease in soluble sugar levels in the MT leaves, as well as the notion that they are being used as an energy source for growth, include an increase in the expression of genes encoding proteins involved in the mitochondrial Tricarboxylic Acid Cycle (TCA) (**Supplementary Fig. S9**). The amino acids produced by protein degradation could fuel the TCA cycle due to the continuous cell wall and vasculature remodelation process.

### MT induces morphoanatomical changes in tomato plants by enhancing protein synthesis and altering essential genes that regulate development, promoting floral transition

The differential transcriptome analysis indicated significant changes in gene expression related to the synthesis of new proteins. In this sense, we found an overrepresentation of genes encoding proteins involved in ribosome biogenesis (*Q*=6.53e^-55^), ribosomal proteins (*Q*=8.2e^-32^), and proteins responsible for translation and processing (*Q*=2.14e^-3^) (**Fig. 3D**). The most differentially ribosomal expressed genes are shown in **Figure 4A**. In this sense, the TOR kinase-regulated pathway promotes growth by facilitating ribosome biogenesis, protein translation and metabolic activation (Dobrenel *et al*., 2016; Shi *et al*., 2018; Van Leene *et al*., 2019; Canal *et al*., 2024). Due to the transcriptional result, we decided to analyse the impact of MT on the TOR pathway by measuring the activation status of the phosphorylated version of the 40S Ribosomal Protein S7 (RPS7-P), a direct target of S7K (Dobrenel *et al*., 2016). We observed an early activation of the TOR pathway three hours after the MT started, compared to plants in normal growth conditions. The TOR-pathway activity levels decreased at six h and remained reduced fifteen days after the treatment ended (**Fig. 4B**), suggesting a rapid activation of this growth-promoting pathway.

**Figure 4.**
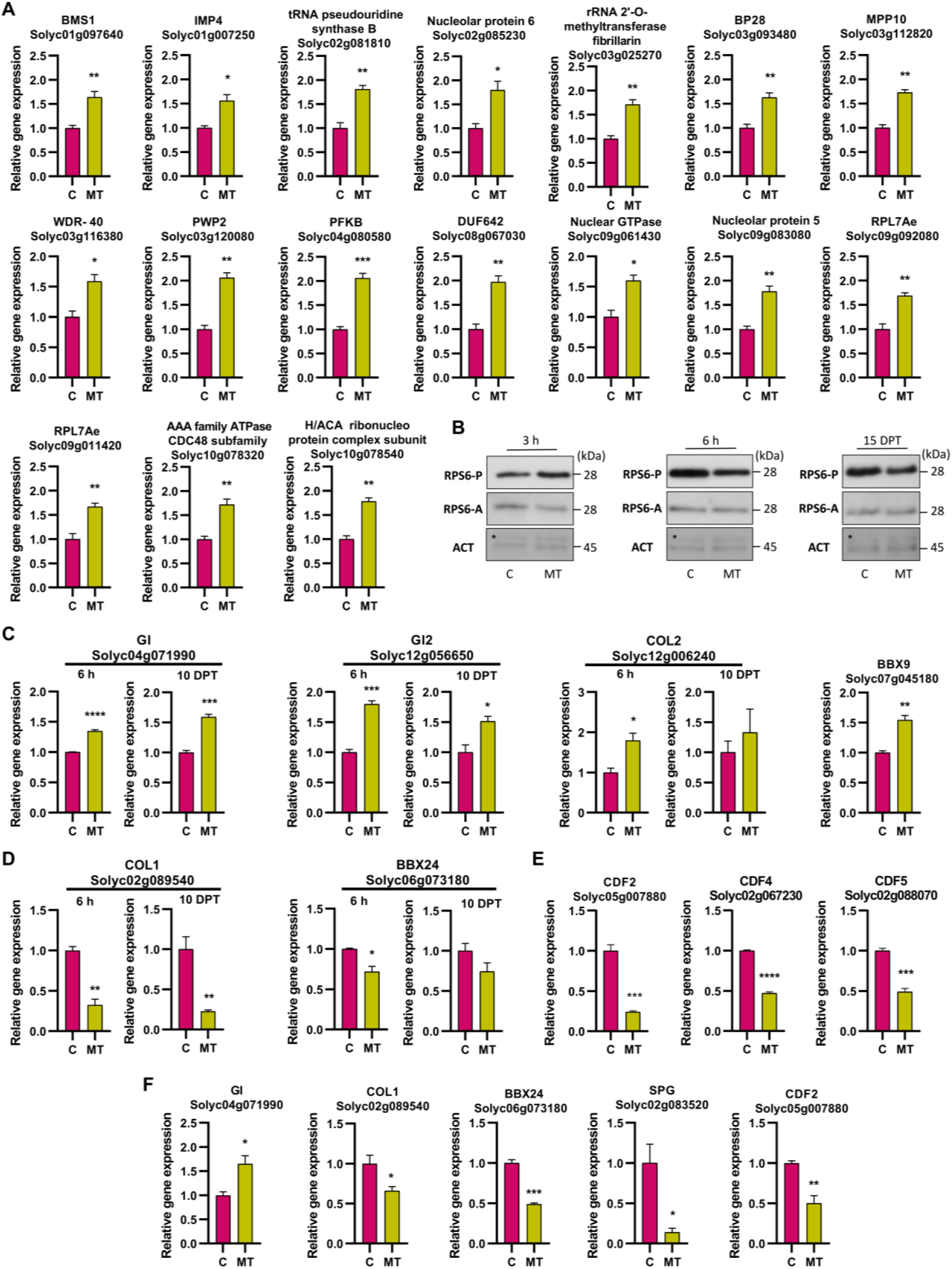
MT induces morphological changes in tomato plants by enhancing protein synthesis machinery and altering developmental genes, thus promoting floral transition. **(A)** The differential expression for genes associated with proteins involved in ribosome biogenesis and function was analysed 6 h after treatment began. **(B)** To assess the activation levels of the TOR pathway, Western blot (WB) analyses were made on total protein extracts prepared from the stem of both control (C) and treated plants (MT) plants at 3 h and 6 h after the start of the treatment, and 15 days post-treatment (DPT). WB membranes were probed with specific antibodies against the phosphorylated version of the 40S ribosomal protein S7 (P-RPS7A, Agrisera # AS194302), total RPS7A (Agrisera #AS194292) and Actin (ACT, Agrisera #AS132640), that was used as a loading control. The (*) indicates a visible band above the actin band, which corresponds to a nonspecific signal generated during the Western blot assay, possibly due to the nonspecific binding of the primary or secondary antibody to another protein in the sample. **(C-F)** Analysis of the expression of critical genes that regulate circadian rhythm and flowering time in tomato plants: **(C)** Genes that show increased expression even after 10 DPT. **(D-E)** Expression levels of genes involved in inflorescence development, specifically those that function as transcriptional repressors, thereby delaying flowering time. **(F)** Expression levels of genes regulating the vegetative and reproductive growth cycle were analysed in floral bud samples from 10-week-old plants. Data are presented as the mean of 3 biological replicates ± SEM, with 3 to 5 plants in each replicate. Samples from control (C) plants are shown in purple, while those from treated (MT) plants are light green. Differences between treatments were considered significant based on an unpaired t-test with a threshold of P < 0.05. Abbreviations: BMS1: BMS1-TYPE G DOMAIN-CONTAINING PROTEIN, IMP4: U3 SMALL NUCLEOLAR RIBONUCLEOPROTEIN PROTEIN IMP4, BP28: U3 SMALL NUCLEOLAR RNA-ASSOCIATED PROTEIN 10, MPP10: U3 SMALL NUCLEOLAR RIBONUCLEOPROTEIN COMPLEX SUBUNIT MPP10P, WDR: WD-40 REPEAT PROTEIN. C-TERMINAL, PWP2: PERIODIC TRYPTOPHAN PROTEIN 2. G-PROTEIN BETA WD-40 REPEAT REGION, PFKB: KINASE PFKB FAMILY CARBOHYDRATE/PURINE KINASE, DUF642: DOMAIN-CONTAINING PROTEIN, RPL7AE: 50S RIBOSOMAL PROTEIN L7AE RIBOSOMAL PROTEIN L7AE/L30E/S12E/GADD45, GI: GIGANTEA, COL: CONSTANS LIKE ZINC FINGER PROTEIN, BBX: B BOX TYPE DOMAIN CONTAINING PROTEIN, CDF: CYCLING DOF FACTOR, SPG: SELF PRUNING.

Tomato plants are day-neutral and have a sympodial growth habit, where the transition to inflorescence occurs at the shoot apices multiple times during its lifespan (Périlleux and Huerga-Fernández, 2022). Synchronising the reproductive phase with the optimal environmental conditions is crucial to maximising the plant’s reproductive potential, partly controlled by the circadian clock (Lee *et al*., 2023). As shown in **Fig. 3C**, genes that regulate the circadian rhythm were overrepresented in MT plants (**Supplementary Fig. S10**). This could be directly linked to the early flowering previously described (**Fig. 2**). Thus, we observed increased expression of genes encoding GIGANTEA (*SlGI*, Solyc04g071990.3 and *SlGI-2*, Solyc12g056650.2) and transcripts for proteins regulated by SlGI, belonging to the MULTIPLE B-BOX (BBX) transcription factors (TF) family (Bu *et al*., 2021; Talar and Kiełbowicz-Matuk, 2021). These included SlBBX7 or CONSTANS-LIKE 2 (*SlCOL2*, Solyc12g006240.2) and SlBBX9 (Solyc07g045180.4) (**Fig. 4C**). Remarkably, the expression of *BBX3* or *CONSTANS-LIKE* 1 (*SlCOL1*, Solyc02g089540.3), and the gene encoding their interacting protein SlBBX24 (Solyc06g073180), was repressed as a consequence of the MT (**Fig. 4D**). SlCOL1 controls inflorescence development by directly binding to the promoter region of the tomato inflorescence-associated gene *SINGLE FLOWER TRUSS (SFT*) and negatively regulating its expression (Cui *et al*., 2022). Suppression of *SlCOL1* leads to the promotion of flower and fruit development, resulting in increased tomato fruit yield (Yang *et al*., 2020; Sun *et al*., 2024). Among the genes observed to be repressed 10 days after the treatment ended, we found several members of the C2C2-DOF (<u>D</u>NA-BINDING WITH <u>O</u>NE <u>F</u>INGER) CYCLIN family, including *SlCDF2* (Solyc05g007880.4), *SlCDF4* (Solyc02g067230.3) and *SlCDF5* (Solyc02g088070.3) (**Fig. 4E**). In this sense, CDF family members act as transcriptional repressors delaying flowering time by regulating FT-like genes (Brandoli *et al*., 2020; Xu *et al*., 2021). Furthermore, while *SlGI* remained induced, the expression levels of *SlCDF2*, *SlCOL1,* and *BBX24* remained diminished in the flower buds of treated plants. At this stage, the expression levels of the *SELF PRUNING* (*SP*), a member of the tomato SP gene family which regulates the vegetative and reproductive growth cycle, acting as a floral inhibitor (Soyk *et al*., 2017) also decreased in MT-plants (**Fig. 4F**).

Among other clock-related genes whose expression is altered after MT, we found those encoding for TIMING OF CAB EXPRESSION 1 (*SlTOC1*, Solyc06g069690.4), whose expression is activated by GI to reset the circadian rhythm (Mizoguchi *et al*., 2005; Mishra and Panigrahi, 2015; Brandoli *et al*., 2020; Liu *et al*., 2024). We also observed increased expression of transcripts encoding proteins from the PSEUDO-RESPONSE REGULATOR or PRR family (*SlPRR3*, Solyc04g049680.2; *SlPRR5*, Solyc03g081240.3; and *SlPRR37*, Solyc04g049670.4) (Irum *et al*., 2024) and a component of the evening complex, EARLY FLOWERING 3 (*SlELF3*, Solyc08g065870.4) (Wu *et al*., 2024) (**Supplementary Fig. S10**). Among the proteins that play a central role in the process of light photoreception, we observed an increase in the expression of transcripts for ELONGATED HYPOCOTYL 5 rs CRYPTOCHROME 2 (SlCRY2, Solyc09g090100.3) and SICRY1b (Solyc12g057040.2). The latter decreases its expression 10 days after the treatment ends compared to the plants used as controls. In tomatoes, SlHY5 influences various developmental processes and light-mediated signalling pathways that affect flowering time and fruit maturation (Wang *et al*., 2021). It acts downstream of photoreceptors and interacts with other signalling components to regulate the timing of flowering in response to light conditions (Zhang *et al*., 2022).

### The effects of MT in tomatoes require an active ethylene pathway

Ethylene alters growth via adjustments in carbon assimilation, central metabolism, and cell wall composition. It influences various developmental processes and physiological responses in plants, including seed germination, cell expansion, flowering, fruit ripening, abscission, and organ senescence (Nascimento *et al*., 2021). Furthermore, a role for the phytohormone ethylene in thigmomorphogenic responses has been proposed (Anten *et al*., 2006). The ethylene biosynthetic pathway involves two specific steps catalysed by ACC (1-AMINO-CYCLOPROPANE-CARBXYLIC ACID) SYNTHASE (ACS) and ACC OXIDASE (ACO). Multigene families encode such enzymes. In tomatoes, 14 *ACS* genes and 7 *ACO* genes have been identified (Liu *et al*., 2015). At 6 h after treatment, we observed an increase in the expression of *ACS 3*, *5 8* and those encoding for ACO2, 3, 4 and 6 (**Fig. 5A**), suggesting a significant increase in ethylene production levels in response to MT. Considering ethylene perception, genes encoding the ETHYLENE RECEPTORS (ETR) ETR 3 to 6 and the EIN3-binding F-box protein 1 (EBF1) were among the group induced after 6 h of MT, some of which were still present 10 days after the end of the treatment (**Fig. 5B**).

**Figure 5:**
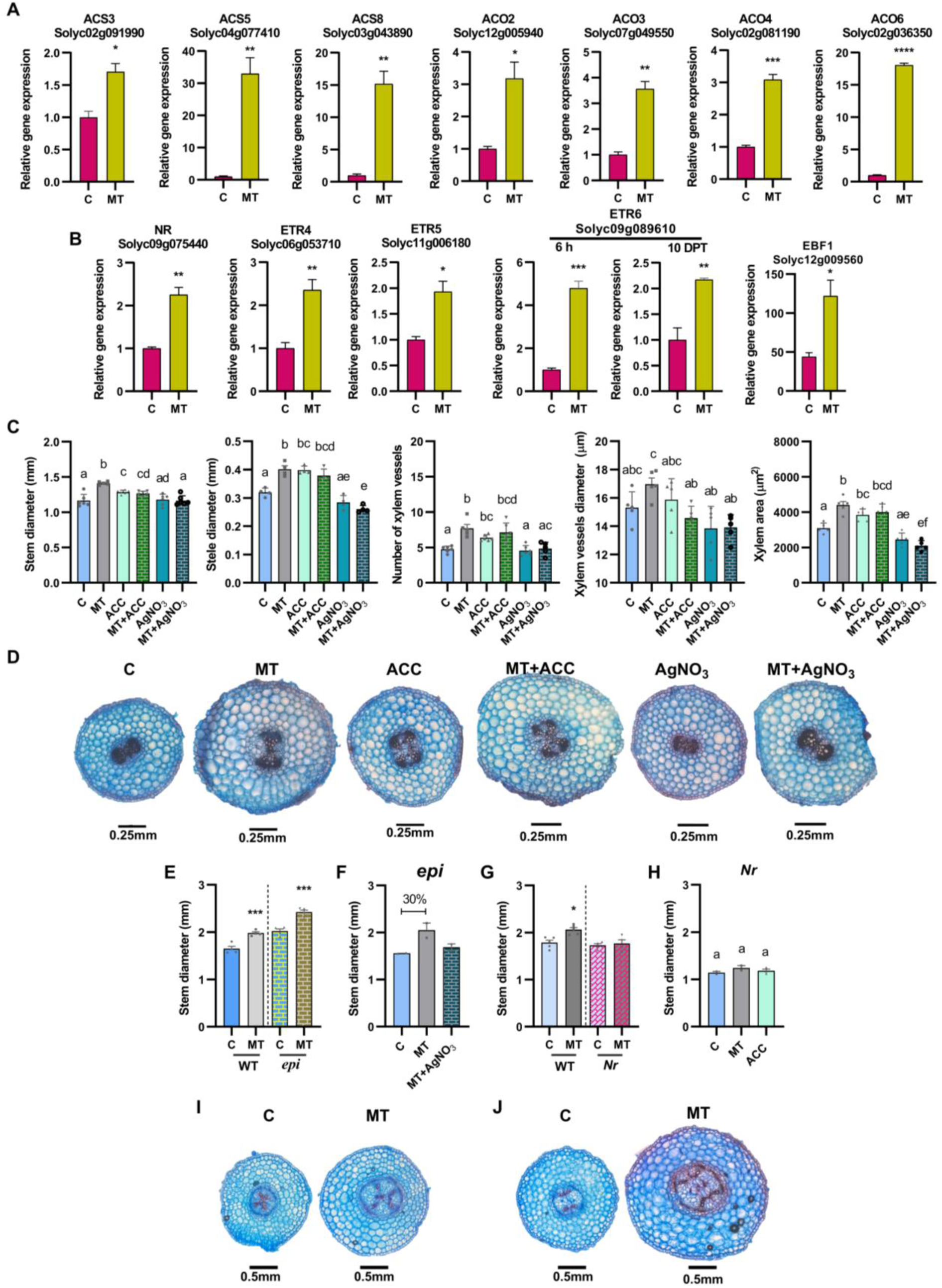
MT triggers an increase in the ethylene pathway, which coordinates the morphoanatomical responses observed. The relative expression of genes involved in **(A)** ethylene biosynthesis and **(B)** perception and signalling was measured at 6 hours and again at 10 days after the initiation of MT treatment. Panels **(C-D)** illustrate the impact of MT on seedlings of AC tomatoes grown *in vitro* for 10 days in vertically arranged square boxes, which were then transplanted to plates under various growth conditions. These conditions included the addition of (1) an ethylene precursor (ACC, 10 µM), (2) an ethylene perception inhibitor (AgNO_3_, 100 µM), and (3) combinations of both with MT (MT+ACC and MT+AgNO_3_). **(C)** Quantitation of parameters such as stem diameter (mm), stele diameter (mm), number of vessels (n), xylem vessels diameter (µm), and total xylem area (µm^2^), performed after 48 h of treatment applied with the 1.5 g device. **(D)** Representative images of stem cross-sections stained with astra-safranin blue after 48 h of 1.5 g-MT, showing the anatomical differences between C and MT tomato seedlings. **(E-F)** Evaluation of the effect of MT on epinastic (*epi*) ethylene-overproducing mutant: **(E)** *epi* seedlings responded to MT by widening their stems at levels comparable to their WT parental plants (VFN8). **(F)** The effect of MT on stem widening in *epi* tomato mutants was abolished in the presence of the ethylene perception inhibitor (AgNO[). **(G-H)** Analysis of Never-ripe (*Nr*) mutants which are insensitive to ethylene: *Nr* mutant plants were unable to respond to **(G)** the MT applied and **(H)** the presence of the ethylene precursor (ACC). **(I-J)** Representative images of stem cross-sections of C- and MT-treated plants of **(I)** WT (VFN8) plants and **(J)** the *epi* mutants. Sections were made from 8 plants per treatment at the end of the MT application. Scale bars represent 0.25 mm and 0.5 mm. Graphs show individual values; data are presented as the mean of 10 plants ± SEM. The figure shows the significant differences between treatments, identified by an ANOVA analysis (Tukey’s test) and unpaired t-test, with a P value < 0.05. The different letters assigned to the groups that present significant variations indicate the differences. Abbreviations: ACS: AMINOCYCLOPROPANE-1-CARBOXYLATE SYNTHASE, ACO: AMINOCYCLOPROPANE-1-CARBOXYLATE OXIDASE, NR: NEVER RIPE. ETR: ETHYLENE RECEPTOR, EBF1: EIN3-BINDING F BOX PROTEIN 1.

Given these observations, we decided to assess the involvement of ethylene in the morphoanatomical responses to MT. AC tomato seedlings were grown *in vitro* for 10 days in square boxes, arranged vertically, in the culture chamber as described in the Methods section (**Supplementary Fig. S1B**). The seedlings were then carefully transplanted to plates with the addition of the ethylene precursor 1-AMINO-CYCLOPROPANE CARBOXYLIC ACID (ACC, 10 µM) or inhibitors of ethylene perception such as silver ions (AgNO_3_, 100 µM) (Zarei and Ehsanpour, 2023). These growth conditions were tested individually (ACC and AgNO_3_) or combined with MT (MT+ACC and MT+AgNO_3_) using 1.5 g weight given the smaller size of the seedlings, always maintaining the treatment time of 48 h. In these different growth conditions, MT also increased stem diameter, mainly contributed by the stele (**Fig. 5C, grey bars**). In addition, an augmented number of xylem vessels was quantified, which, although not significantly larger in diameter, led to a greater xylem vascular area (**Fig. 5C, grey bars**). Surprisingly, a similar effect was observed after 10 µM ACC addition without MT (**Fig. 5C, light green bars**). The combined MT plus the addition of the precursor (MT+ACC) did not produce an additive effect; the observed changes were comparable to those generated by MT or ACC separately (**Fig. 5C, green bar**). This non-additive effect may be due to stem cell growth, expansion limitations, or treatment time. The most remarkable observation was that the addition of an ethylene inhibitor (silver ions) in combination with mechanical treatment (MT + AgNO_3_) completely abolished the impact of MT on stem outgrowth and all associated parameters (**Fig. 5C, dark blue bars**), demonstrating that an active ethylene signalling pathway is necessary to generate the morphoanatomical changes. The quantifications were made by performing histological sections, as shown with the representative stem cross-sections in **Fig. 5D**.

To further investigate ethylene involvement in MT responses, we examined the impact of MT on *epinastic* (*epi*) mutants that are ethylene overproducers, showing severe leaf epinasty, thickened stems and petioles, and a compact growth habit (Barry *et al*., 2001). They exhibited thicker stems under normal conditions and, consistently with our experimental design, were able to respond to MT at levels comparable to their WT parental plants (cv. VFN8) (**Fig. 5E**), as long as the ethylene perception was not inhibited (**Fig. 5F**). This result can be observed in the histological sections of stems from plants subjected to MT, of wild type WT (VFN8) (**Fig. 5I**) and the mutant (*epi*) (**Fig. 5J**). We then analysed *Never ripe (Nr)* mutants, which carry a mutation in the ethylene receptor gene (*SlETR3*, Solyc09g075440), conferring a reduced ethylene sensitivity (Lanahan et al., 1994). We found that *Nr* plants could not widen their stems by MT (**Fig. 5G**) nor by the exogenously added ethylene in the form of ACC (**Fig. 5H**). These results confirmed that an active ethylene signalling pathway was necessary to achieve the morpho-anatomical changes induced in the stem due to MT.

### Auxins are involved in the mechanism of MT-induced effect and are necessary for ethylene action

Auxins play critical roles in developing vasculature in tomato plants, modulating vascular tissue formation, and affecting cell division and elongation (Petrás□ek and Friml, 2009). They bind to receptor proteins on the cell membrane, activating ATPase proton pumps to acidify the cell wall and soften it by increasing internal pressure. XYLOGLUCAN ENDOTRANSGLYCOSYLASES/HYDROLASES (XTHs) and EXPANSINS (EXPs) modify cell wall properties and promote cell expansion (Perrot Rechenmann, 2010). In our transcriptome, we observed that MT notably upregulated certain expansins and β-glucanases and some XTH gene family members after 6 hours of MT (*EXPANSIN 2* (Solyc06g049050), *EXPANSIN 12* (Solyc05g007830), *EXPANSIN S2* (Solyc07g063210) and *EXPANSIN A11* (Solyc04g081870)). Additionally, we noted that several XTHs and other genes encoding modifying enzymes like *XTH8* (Solyc07g006870), *XTH5* (Solyc11g040140), *GLUCAN SYNTHASE LIKE 1* (Solyc07g0539809), and *GLUCAN SYNTHASE LIKE 3* (Solyc07g061920) remained upregulated after 10 days of MT.

Therefore, we decided to examine the influence of auxins on the phenotypic changes observed in tomato plantain response to MT. We analysed the effect on stem width of adding exogenous auxins, such as INDOLE-3-ACETIC ACID (IAA, 3-IAA), or inhibiting polar transport using N-1-NAPHTHYLPHTHALAMIC ACID (NPA), alone and in combination with MT. We used plate-grown AC tomato seedlings for 10 days and exposed them to different conditions. First, we observed that 1.5 g-MT increased the stem diameter, mainly due to the enlargement of the stele, which led to a more significant number of xylem vessels and an increased xylem vascular area (**Fig. 6A, grey bars**).

**Figure 6.**
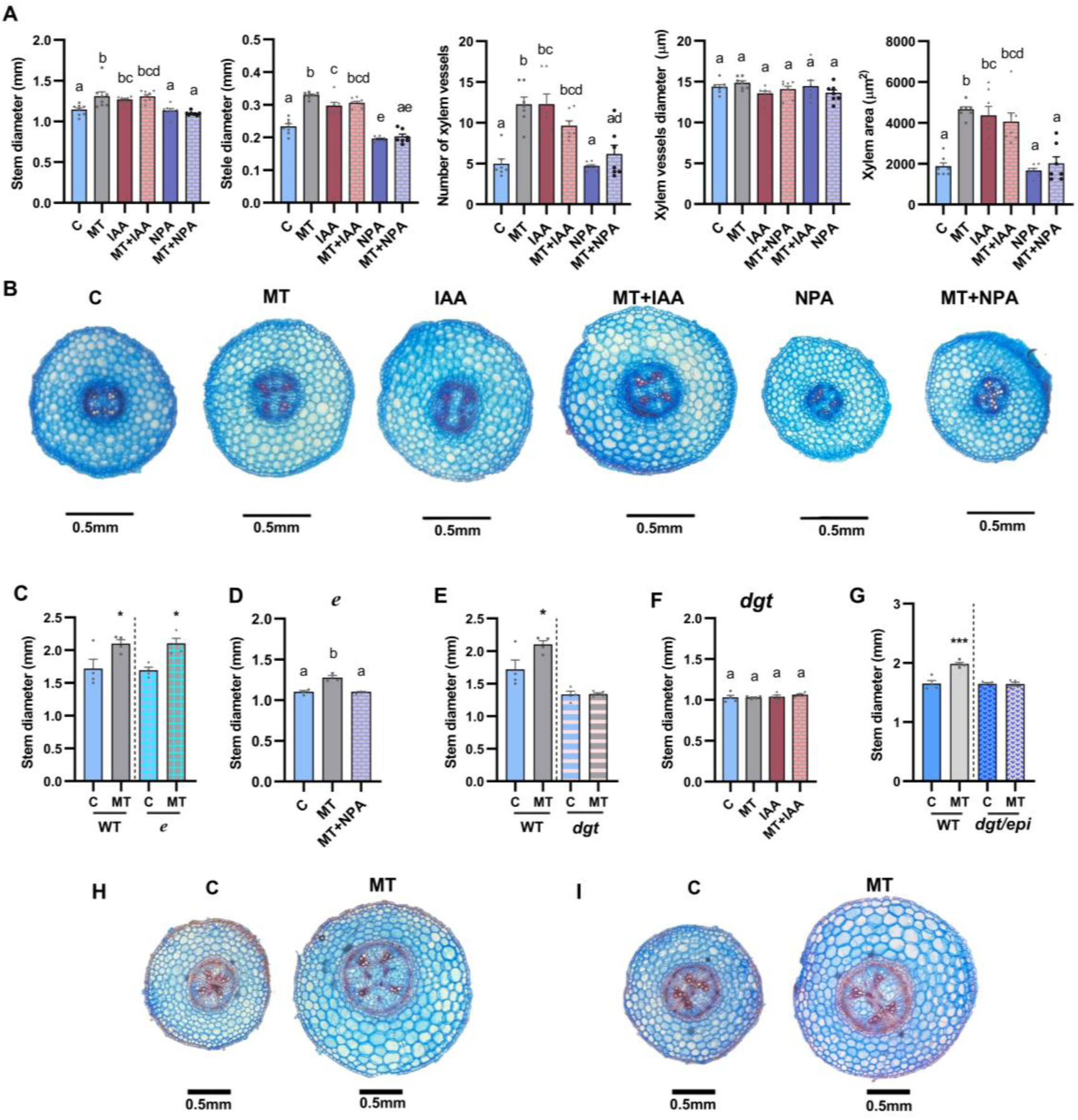
MT induces response mechanisms that involve the action of auxins. **(A)** Analysis of the impact of adding an auxin precursor (IAA, 1 µM) or a polar transport inhibitor (NPA, 1 µM), both individually and in combination with a treatment using a 1.5 g device (MT+IAA and MT+NPA). Quantitation of parameters such as stem diameter (mm), stele diameter (mm), number of vessels (n), xylem vessels diameter (µm), and total xylem area (µm^2^), performed on 10-day-old AC seedlings after 48h of treatment. **(B)** Representative images of stem cross-sections stained with astra-safranin blue after 48 h of 1.5 g MT, showing the anatomical differences between C and MT tomato seedlings. **(C)** Impact of MT on the stem diameter widening in the auxin-pathway mutant *entire* (*e*), which exhibits an enhanced response to auxin, compared to WT-AC tomato seedlings. **(D)** Application of NPA abolished the effect of MT on stem diameter enlargement in the WT and *e* mutants. **(E)** The *diageotropic* (*dgt*) tomato mutant, characterised by a reduced sensitivity to auxins, did not show stem enlargement in response to MT. **(F)** The addition of IAA did not affect the stem widening in the *dgt* mutant. **(G)** Analysis of the stem enlargement response to MT in the double mutant (*epi*/*dgt*). No response to MT was observed in *epi*/*dgt* double mutant compared to WT tomato seedlings. **(H, I)** Representative images of stem cross-sections from control and MT-treated plants, corresponding to **(H)** WT (AC) tomatoes and **(I)** *e* mutant plants. Cross-sections were made randomly from 5 plants per treatment at the end of the MT application. Scale bars represent 0.5 mm. Graphs show individual values; data are presented as the mean of 10 plants ± SEM. Differences between treatments were considered significant according to multiple comparisons ANOVA (Tukey’s test) and unpaired t-test, with a P value < 0.05.

**Figure 7.**
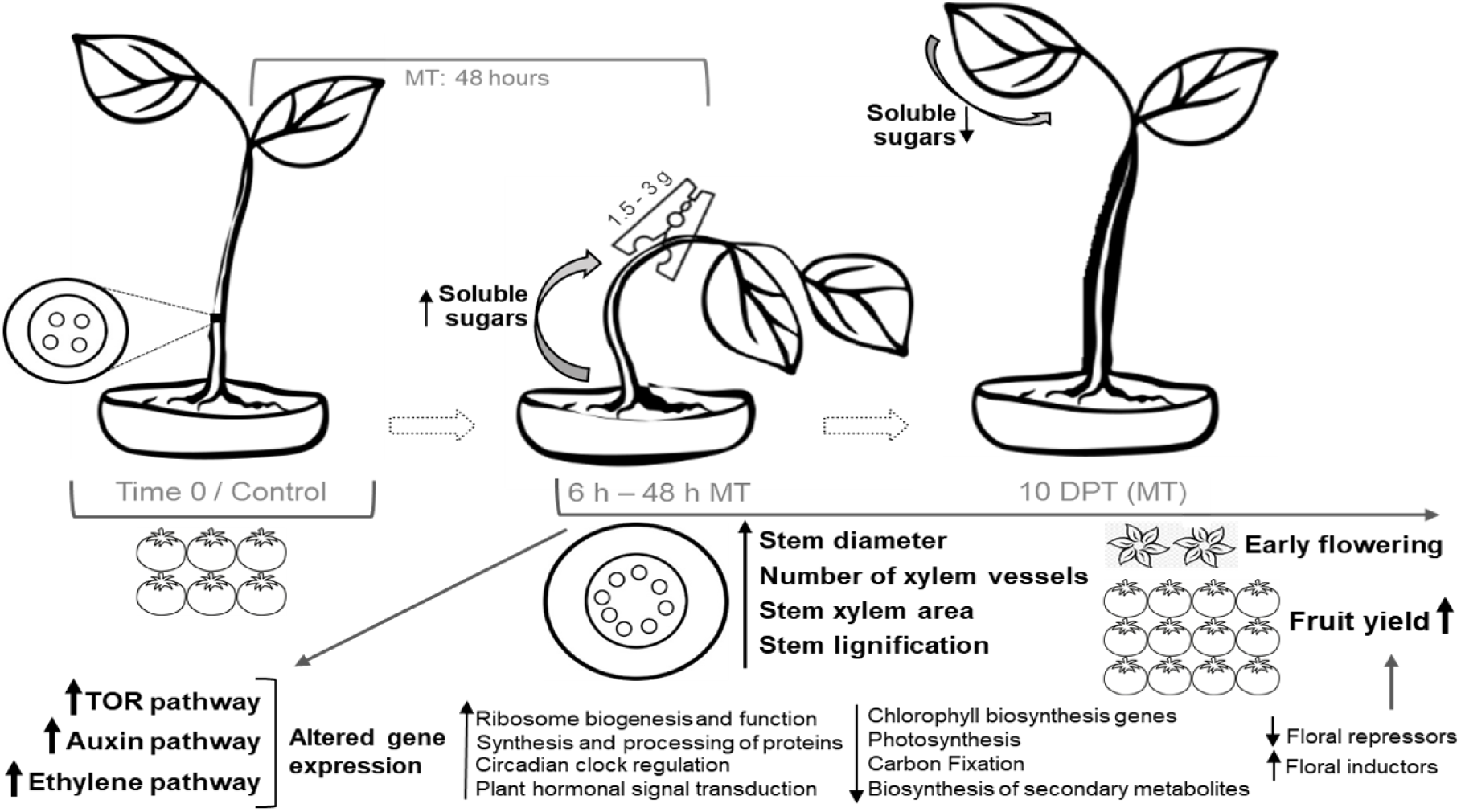
Schematic representation of the morphological and physiological impacts triggered by combined MS in tomato plants. Ten–day–old tomato plants exposed to MT with a variable weight of 1.5 to 3 g, positioned on the upper part of the stem for 48 h, exhibited multiple layered responses. Initially, this exposure significantly increased stem width, stele diameter, and the number of xylem vessels, leading to a substantial increase in stem xylem area. All the changes were observed as early as 6 hours after the initiation of MT and continued until nearly 10 DPT. MS also impacted the lignification of the stem tissues. MT induced changes in gene expression and altered plant metabolism to sustain the morphological and physiological changes, amplifying the initial responses throughout the entire life cycle of the plants. As a result, there was an increase in the activation of the Target of Rapamycin (TOR) pathway, along with the induction of the auxin and ethylene pathways, which are hormonal pathways required for the observed phenotypic responses. All these changes impacted plant development, leading to early flowering and increased fruit yield in the tomato plants subjected to MT. The up arrow means an increment or up-regulation, and the down arrow represents a decrease or down-regulation.

Surprisingly, a similar effect was observed with the addition of IAA 1 µM alone (**Fig. 6A, red bars**) or in combination with MT (MT+IAA) (**Fig. 6A, pink bars**). These two conditions had no additive effect (**Fig. 6A, pink bars**). Inhibition of auxin distribution by the addition of NPA 1 µM alone (**Fig. 6A, purple bars**) or in combination with MT (MT+NPA) (**Fig. 6A, light purple bars**) completely annulated the effect of MT on stem width enlargement, resulting in a phenotype comparable to that of control (C) plants **(Fig. 6A, light blue bars**). The quantifications were made by performing histological sections, as shown with the representative stem cross-sections in **Fig. 6B**.

Then, we explored the possible connections in depth using plants with alterations in the auxin pathway. We evaluated the impact of MT on *entire* (*e*) plants that have mutated a transcriptional repressor of auxin signalling (*AUX/IAA9*) and thus have an increased response to hormone addition (Zhang *et al*., 2007). We also tested the *diageotropic* (*dgt*) mutant, which has a mutated cyclophilin biosynthesis allele, reducing sensitivity to auxins (Oh *et al*., 2006). We observed that *e* plants responded to MT by widening their stems (**Fig. 6C, H, I**), and such effect was lost in the presence of NPA (**Fig. 6D**). In contrast, *dgt* mutants were unable to enlarge their stems, even when exogenous IAA was applied (**Fig. 6F**). This last observation further helped us confirm the mutant identity and confirmed that auxins are necessary for tomato plant responses to the MS stimuli induced by the treatment. Finally, we evaluated the ethylene and auxin pathways using the double mutant *epi/dgt*, which overproduces ethylene but is less sensitive to auxin. We observed that the *epi/dgt* plants could not respond to MT (**Fig. 6G**), reinforcing that an active auxin pathway is necessary for ethylene to effectively influence tomato widening and the increment of the vascular area.

## Discussion

Plants often experience mechanical forces from increasing body weight and surrounding cell growth and expansion. Developmental plasticity allows them to modulate their phenotype depending on environmental conditions, influencing biomass and seed yield efficiency. (Sultan, 2010; Du and Jiao, 2020; Sampathkumar, 2020). The exogenous factors causing MS due to MT impacting on the stem can be classified based on how they are applied, as those induced by gravity (i.e. snow), by gradients in pressure, or caused by ‘thigmo stimuli’ which lead to thigmomorphogenesis (Telewski, 2016; Kouhen *et al*., 2023; Khan *et al*., 2024).

Our research devised a combined treatment on tomato seedlings, modifying previous well-documented studies conducted on adult Arabidopsis and sunflower plants that influenced the secondary growth, resulting in changes in stem thickness and the development of vascular vessels (Ko *et al*., 2004; Cabello and Chan, 2019). Remarkably, we did not observe any changes in the height of the tomato plants, which contrasts with the expected classical thigmomorphogenetic reported responses (Johjima *et al*., 1992; Coutand, 2000; Van Gaal and Erwin, 2005, Saidi *et al*., 2009; Wang, 2024). Other forces may also affect the plants, such as increasing stem diameter. The bending caused by the weight could create conditions similar to those found in microgravity combined with hypergravity phenomena (Kouhen *et al*., 2023). Microgravity typically promotes elongation and suppresses lateral expansion in shoot organs, whereas hypergravity inhibits elongation and enhances lateral expansion. In Arabidopsis, this response is primarily influenced by the orientation of cortical microtubules and is regulated by the 65 kDa MICROTUBULE-ASSOCIATED PROTEIN-1 (MAP65-1) (Soga *et al*., 2018). Our transcriptional data indicated that the homologue MAP65-1 in tomato (Solyc01g005080.3) and other genes encoding proteins associated with microtubule orientation were downregulated in response to our MT. This suggests that our plants exhibit a combined response due to microtubule rearrangements caused by the MT (Ghosh *et al*., 2021; Jonsson *et al*., 2022).

Ethylene and auxins are key players in regulating vascular development and secondary growth (Brenya *et al*., 2022; Wang, 2024). Ethylene influences plant growth, including photosynthesis and leaf senescence (Ceusters and Van de Poel, 2018; Mohorović *et al*., 2024). Ethylene overproducer plants show enhanced vascular cell division and increased vascular size in the stem (Etchells *et al*., 2012; Khan *et al*., 2024;). Ethylene, produced in response to environmental cues such as leaning, stimulates cell division in the cambial meristem, forming wood (Love *et al*., 2009). It supports body-weight-induced secondary growth in Arabidopsis, highlighting its importance in vascular development. In this sense, we demonstrated that ethylene influences and an intact ethylene pathway are needed for the plant responses to our imposed MT (**Fig. 5**). In our experiment, the increment of ethylene levels could be included in the physiological levels, as we only detected downregulation of chlorophyll synthesis and photosynthesis genes (**Supplementary Fig. S5 and S7**), consistent with the changes described recently by Mohorović and colleagues (2024). The repression of photosynthesis-related genes without impact on the photosynthesis rate was also described for transgenic plants expressing the sunflower gene HaHB4 (Manavella *et al*., 2008), reinforcing the continuous crosstalk between ethylene signalling and photosynthesis. However, we found no significant alterations in chlorophyll and starch content after MT (**Supplementary Fig. S6**) and only observed a reduction in the flowering time (**Fig. 2**).

We proposed that the downward bending of the stem due to added weight causes the leaves to bend downwards, resembling an “epinastic” phenotype, reducing the leaf’s coverage and ability to form a dense canopy. As a result, the light interception was affected, which hindered the photosynthesis process and decreased the expression of photosynthetic machinery and chlorophyll synthesis (**Supplementary Fig. S5 and S7**). In addition, the CO_2_ fixation through the Calvin cycle decreased (**Supplementary Fig. S8**), perhaps because of the lowering in some of the substrates that stimulate this cycle, such as NADPH and ATP, which come from the light reactions of the thylakoid membranes. Overall, we observed a reduced expression of the genes coding for proteins belonging to the photosynthetic machinery, the light-harvesting reactions, carbon fixation and regeneration in a similar weight to those observed in plants exposed to ethylene (Nascimento *et al*., 2021; Mohorović *et al*., 2024) or with higher ethylene content (Manavella *et al*., 2008). Thus, the metabolic effects and gene expression changes may stem from reduced light interception and increased ethylene production. Research showed that low light stress can downregulate Rubisco subunit genes, decreasing carboxylation activity (Sun *et al*., 2014; Liu *et al*., 2020). Ethylene inhibits net carbon exchange, but repositioning leaves toward light can reverse this effect, indicating that ethylene indirectly impacts carbon gain by reducing light capture through epinasty (Woodrow *et al*., 1989).

We included the role of auxins as hormones responsible for the impact of the stem widening and generation of vascular bundles. Auxins play a crucial role in stem widening and vascular bundle formation in tomato plants, regulating vascular tissue development (**Fig. 6**). Their distribution is essential for an efficient and functional vascular system, stimulating cell division and cambium activity, which leads to xylem and phloem development for nutrient transport (Tuominen *et al*., 1997; Nilsson *et al*., 2008; Petrás[ek and Friml, 2009). In this sense, it has been shown that applying weight to the main stem of Arabidopsis plants induces cambial differentiation, enhancing auxin transport and promoting secondary xylem development (Ko *et al*., 2004; Cabello and Chan, 2019). Epinasty can be caused by ethylene and high auxin concentrations.

Floral transition in tomatoes thus marks the switch of the SAM from a monopodial program to a sympodial patterning characterised by the formation of shoots and inflorescence. Our study on combined MS observed a phenotypic reduction in flowering time, correlating with the transcriptional changes observed (**Fig. 2**). In this sense, it is well known that GI activates the expression of *CO* and *SFT* by facilitating the degradation of transcriptional repressors belonging to the CDF protein family (Brandoli *et al*., 2020; Yang *et al*., 2020).

Controlling flowering time improves crop yield (Rajendran *et al*., 2021). Synchronising the reproductive phase with the optimal environmental conditions is crucial to maximising the plant’s reproductive potential, partly controlled by the circadian clock (Lee *et al*., 2023). As shown (**Fig. 3**), genes that regulate the circadian rhythm were overrepresented in MT-plants (**Supplementary Fig. 10**).

As the global population continues to grow, the challenge of climate change makes it increasingly urgent to find ways to increase crop production. Our study demonstrates that combined MT is an environmentally friendly approach and holds immense potential to stimulate plant growth, offering a promising solution for the future of sustainable agriculture (Kouhen *et al*., 2023)

In our research, we aimed to investigate the influence of hormones such as ethylene and auxins on the development of vascular tissues in tomatoes. We also sought to understand the relationship between these hormones and the morphological changes in the stem and how these changes can ultimately impact plant production. We focused on identifying sustainable strategies to enhance crop yield (Ghosh *et al*., 2021). To achieve this, we designed a successful experiment combining MT to induce MS responses in tomato plants, promoting increased crop production. We speculated that prolonged MT could program somatic memory, priming for defence acclimation and creating solutions to improve agricultural sustainability.

(Ghosh *et al*., 2021; Brenya *et al*., 2022).

## Supplementary data

**Supplementary Figure S1. Illustrative pictures and schemes of the systems used for applying mechanical treatment (MT) on tomato plants**

**(A)** Application of the wooden clip-type device placed on the stem of 4-5 cm tall seedlings (approximately 10-day-old plants) grown in pots. **(B)** Applying the MT on the stem apex of seedlings cultivated in vitro in vertically arranged square plates containing MS1X, agar 1 %. **(C)** Measure the stem diameter at the end of the treatment, approximately 1-2 cm from the base, using a digital calliper. See Supplementary Video S1 for details.

**Supplementary Figure S2. The impact of MT on stem morphoanatomy is evident within the first six hours after the start of treatment**

Analysis of stele diameter (mm), number of vessels (n), xylem vessels diameter (µm), and total xylem area (µm^2^), performed in stem cross-sections of seedlings subjected to MT (grey bar) compared to control plants (C) (light blue bar). Evaluations were performed at the following time points: **(A)** 3 h, **(B)** 6 h, **(C)** 12 h, **(D)** 24 h and **(E)** 48 h after the initiation of treatment, consisting of the application of 3 g device as described in Material and Methods (see Supplementary Video S1). Images on the right correspond to representative histological sections of stems stained with astra-safranin blue double staining. Scale bars represent 0.5 mm. Graphs show individual values; Data are presented as the mean of 6 plants ± SEM. The figure shows the significant differences between treatments, identified by an ANOVA analysis (Tukey’s test). The differences are indicated with different letters assigned to the groups that present significant variations and asterisks indicate significant differences between control (C) and treated (MT) plants, according to an unpaired t-test at P < 0.05.

**Supplementary Figure S3. The impact of MT on stem widening is observed in different tomato varieties**

**(A-C)** Response of the Ailsa Craig (AC), M82, and Money Maker (MM) tomato varieties to treatment (MT) with a 3 g device applied to the stem apex of 10-day-old plants for 48 h. **(D-E)** Effect of treatment with 1.5 g device MT on the stem of 10-day-old plants of the Chalchalero®, Platense and Regina® commercial varieties. Graphs show individual values; data are presented as the mean of 8 plants ± SEM. Control (C) plants are represented in light blue bars, while treated (MT) plants are represented in grey bars. Measurements were taken at two time points: “Zero hours” (O h), which indicates the stem diameter before starting the treatment, and “48 h,” marking the end of the MT period. Significant differences between treatments and the control are indicated by different letters, according to the ANOVA test with multiple comparisons (Tukey’s test), for a significance level of P < 0.05.

**Supplementary Figure S4. The impact of MT on the production of commercial tomato Chalchalero®**

**(A)** The impact of MT on stem width (mm) was observed 48 h post-treatment. **(B)** Number of fruit and **(C)** weight (g) of fruits collected from plants aged 28 to 38 weeks after sowing (WAS) in the MT group (light green) compared to the control group (C) (pink). **(D)** The total number of fruits harvested at the end of the life cycle is shown in C (pink bars) and MT (light green bars) plants. **(E)** Comparison of the total fruit yield (g/plant) obtained from C and MT plants by the end of their life cycle. **(F)** Representative images show the total tomato fruits collected at 28 WAS in control (C) and treated (MT) plants. The graphs display individual values, with means representing the average of 10 plants ± SEM. Significant differences between treatments and the control are indicated with different letters, according to the ANOVA analysis with multiple comparisons (Tukey’s test). Likewise, differences between treatments are considered significant if determined by a t-test with a P value < 0.05.

**Supplementary Figure S5. Expression level of genes involved in chlorophyll biosynthesis and catabolism**

Schematic representations of **(A)** the chlorophyll biosynthesis pathway and **(B)** the degradation pathway, adapted from the KEGG PATHWAY Database (Okuda *et al*., 2008). Different coloured boxes illustrate data based on gene regulation and their level of statistical significance. Blue boxes represent down-regulated genes, violet boxes indicate up-regulated genes, and the light blue box in the middle of the reference scale corresponds to genes with non-significant differences. Within each colour category, the intensity varies according to the P value (*P < 0.05; **P < 0.01, ***P < 0.001), with greater colour intensity for higher levels of statistical significance. **(C)** The impact of MT on the expression of genes associated with chlorophyll biosynthesis was observed 6 h after the start of the MT. **(D)** The altered expression of genes that regulate chlorophyll degradation is influenced by the effect of MT after 6 h of treatment, with low expression levels maintained for up to 10 days post-treatment. Graphs were generated from RNA-seq data using the FRAGMENTS PER KILOBASE OF TRANSCRIPT PER MILLION MAPPED READS (FPKM) normalisation method. Graphs represent the mean of 3 biological replicates ± SEM. Differences between control and mechanical treatment (MT) groups were assessed using an unpaired t-test and were considered significant when P < 0.05. Abbreviations: MPEC: MAGNESIO PROTOPORFIRINA IX MONOMETIL ÉSTER CICLASA, DVR: ACTIVIDAD 3,8-DIVINIL PROTOCLOROFILIDA A 8-VINIL REDUCTASA, PORC: PROTOCLOROFILIDA OXIDORREDUCTASA, POR: PROTOCLOROFILIDA OXIDORREDUCTASA, NOL: CLOROFILA B REDUCTASA, NYC1: CHLOROPHYLLIDE B REDUCTASE NYC1 CHLOROPLASTIC.

**Supplementary Figure S6. Total chlorophyll and starch content in leaves and stems of tomato plants subjected to MT**

**(A)** Total chlorophyll content (µg/mg) and **(B)** starch quantification (nmol/mg) at 24 h, 48 h after the application of MT and 10 days post-treatment, measured in leaves and stem of Control (C, violet bars) and treated (MT, yellow bars) plants. The values represent the men of 3 plants ± SEM. There were no significant differences between treatments according to the unpaired t-test with a P value < 0.05.

**Supplementary Figure S7. Expression levels of genes involved in the photosynthetic machinery and antenna proteins**

Expression levels of genes associated with **(A)** photosystem II, **(B)** photosystem I, **(C)** those associated with electron transport in the photosynthetic machinery, **(D)** F-type ATPase subunit members and **(E)** the chlorophyll a/b-binding protein genes, evaluated in plants subjected to MT (light green) compared to control conditions (purple). Repression of genes was observed 6 h after the start of MT, and some of them remained repressed even 10 days after treatment. The data were obtained using the FRAGMENTS PER KILOBASE OF TRANSCRIPT PER MILLION MAPPED READS (FPKM) normalisation method from RNA-Seq analysis. The graphs represent the mean of 3 biological replicates ± SEM. Differences were considered significant according to the unpaired t-test with a P-value (*P < 0.05; **P < 0.01). Abbreviations: PSBO: OXYGEN EVOLVING ENHANCER PROTEIN I, CHLOROPLASTIC. PSBQ: PHOTOSYSTEM II OXYGEN EVOLVING COMPLEX PROTEIN III, PSBR: PHOTOSYSTEM II 10 KDA POLYPEPTIDE, CHLOROPLASTIC, PSBS: PHOTOSYSTEM II 22 KDA PROTEIN, CHLOROPLASTIC, PSBW: PHOTOSYSTEM II PROTEIN CLASS II. PSBX: PHOTOSYSTEM II 23 KDA PROTEIN, PSBY: PHOTOSYSTEM II CORE COMPLEX PROTEINS, CHLOROPLASTIC, PSAD: PHOTOSYSTEM I REACTION CENTER PROTEIN SUBUNIT II, CHLOROPLASTIC. PSAE: PHOTOSYSTEM I REACTION CENTER SUBUNIT IV A, PSAF: PHOTOSYSTEM I REACTION CENTER PROTEIN, SUBUNIT III, PSAG: PHOTOSYSTEM I REACTION CENTER SUBUNIT V, CHLOROPLASTIC, PSAH: PHOTOSYSTEM I REACTION CENTER SUBUNIT VI, PSAK: PHOTOSYSTEM I REACTION CENTER SUBUNIT X, PSAL: PHOTOSYSTEM I REACTION CENTER SUBUNIT XI, PSAN: PHOTOSYSTEM I REACTION CENTER SUBUNIT N, CHLOROPLASTIC, PC: PLASTOCYANIN, CHLOROPLASTIC, FNR: FERREDOXIN NADP REDUCTASE, CHLOROPLASTIC. ATPC: ATP SYNTHASE GAMMA CHAIN CHLOROPLASTIC, ATPD: ATP SYNTHASE DELTA CHAIN CHLOROPLASTIC, CAB: CHLOROPHYLL A-B BINDING PROTEIN CHLOROPLASTIC.

**Supplementary Figure S8. Expression of genes involved in the Carboxylation and Sugar metabolism pathway**

**(A)** Schematic representation of the carbon fixation pathway and plant sugar metabolism, modified from the KEEG pathway database. **(B)** The relative expression levels of genes associated with carbon fixation and reduction during the Calvin cycle are depicted. A significant decrease in the expression of genes related to carbon fixation, such as *RBCS1*, *RBCS2*, and *RBCS5*, was observed 6 hs after the treatment (MT). This decline was also noted in genes involved in RuBP regeneration (such as RCA) and sugar reduction (including PGK-1 and GAPDH), with some cases showing persistence of this decrease up to 10 days post-treatment. **(C)** The expression level of genes involved in sugar synthesis, such as FBA and FBP, was reduced during the glycolysis stage. The diagram illustrates the data by different coloured boxes according to gene regulation and statistical significance: blue boxes represent down-regulated genes, violet boxes represent up-regulated genes, and the light blue boxes correspond to genes with non-significant differences. Within each colour category, hue varies as a function of P value (*P < 0.05; **P < 0.01, ***P < 0.001), with greater colour intensity for higher levels of statistical significance. Data are presented as the mean of 3 biological replicates ± SEM. Differences were considered significant by unpaired t-test at P < 0.05. Abbreviations: RBCS: RIBULOSE BISPHOSPHATE CARBOXYLASE SMALL SUBUNIT CHLOROPLASTIC, RuBP: RIBULOSE-1,5-BISPHOSPHATE, PGK-1: PHOSPHOGLYCERATE KINASE 1, RCA: RIBULOSE BISPHOSPHATE CARBOXYLASE/OXYGENASE ACTIVASE CHLOROPLASTIC, GAPDH: GLYCERALDEHYDE 3 PHOSPHATE DEHYDROGENASE, FBA: FRUCTOSE BISPHOSPHATE ALDOLASE, FBP: FRUCTOSE 1,6 BISPHOSPHATASE, CYTOSOLIC.

**Supplementary Figure S9. Expression level of genes involved in Glycolysis and the Tricarboxylic Acid (TCA) cycle**

**(A)** Diagram of the metabolic flux of glycolysis and the TCA cycle, highlighting the enzymes encoded by the analysed genes. **(B)** Gene expression levels of key enzymes involved in glycolysis, such as PFK and PK, showed a significant decrease 6 h after initiation of MT. The expression of GAPDH remained repressed even 10 days post-treatment. **(C)** Expression of genes associated with the TCA cycle and GABA synthesis observed increased at 6h 6 hours after the MT began. Data were obtained using the fragment normalisation method per kilobase of transcript per million mapped reads (FPKM) from RNA-Seq analysis. Graphs represent the mean of 3 biological replicates ± SEM. Differences were considered significant by unpaired t-test at P < 0.05. Abbreviations: GAPDH: GLYCERALDEHYDE 3 PHOSPHATE DEHYDROGENASE, PFK: ATP DEPENDENT 6 PHOSPHOFRUCTOKINASE, PK: PYRUVATE KINASE. CS: CITRATE SYNTHASE, KAT: 3 KETOACYL COA THIOLASE 2, SUCLA: SUCCINYL COA LIGASE BETA SUBUNIT, MITOCHONDRIAL, SUCDH: SUCCINATE DEHYDROGENASE MITOCHONDRIAL, FH: FUMARATE HYDRATASE, MDH: MALATE DEHYDROGENASE, SCOA: SUCCINYL COA LIGASE ALPHA 1 SUBUNIT, GAD: GLUTAMATE DECARBOXYLASE, ACO: ACONITATE HYDRATASE, GABA-TP: GAMMA AMINOBUTYRATE TRANSAMINASE ISOFORM.

**Supplementary Figure S10. Effects of MT on regulating genes associated with the circadian cycle and light perception**

**(A)** Increased expression levels of genes related to the circadian clock at 6 h after treatment. **(B)** Expression of circadian clock-related genes remained elevated 10 days post-treatment. **(C)** The expression of specific genes associated with the circadian cycle was measured in flower buds from plants at approximately 10 weeks old. **(D)** The expression of a gene related to light perception was reduced in the flower buds. Tubulin was employed as the reference gene for real-time RT-qPCR analyses. Data are presented as the mean of three plants ± SEM. Differences between treatments were considered significant based on an unpaired t-test at P < 0.05. Abbreviations: ELF3: EARLY FLOWERING 3, PRR: PSEUDO-RESPONSE REGULATORS, TOC1: (TIMING OF CAB EXPRESSION 1), GI: GIGANTEA, HY5: ELONGATED HYPOCOTYL 5.

**Supplementary Table S1.** Oligonucleotides used in this study.

**Supplementary Video S1.** The use of MT as an eco-friendly technology for enhancing tomato production.

## Abbreviations

MS: Mechanical stress
MT: mechanical treatment

## Acknowledgements

We thank Dr María Laura Vidoz (Plant Physiology Laboratory, FCA-UNNE, Instituto de Botánica del Nordeste, IBONE-CONICET) for sharing tomato mutant plants in hormonal pathways. The authors thank Dr Paula Filippone and Dr Ana Ramallo for their helpful discussions, participation in outreach lectures, and the work that will be included in future research. We also thank Dr Patricia L. Albornoz (FCNe IML, Universidad Nacional de Tucumán and Fundación Miguel Lillo) for her valuable technical assistance in teaching us to perform histological sections. Thank Mg, Elena Fanny Jerez, for collaborating on the fruit quality analysis (EEA Famaillá, INTA). We are particularly grateful to Mg. Victoria Castro Demiryi for her enthusiastic support in documenting with videos showing MT treatments and Dr Matías Capella for his participation in them.

## Author contributions

JC-E performed most of the experiments with AC variety, analysed data and prepared the figures. SMS and JAMM performed the experiments with the commercial varieties and determined tomato quality. JVC participated in the design of the original idea and discussed the results. RLC and EW conceived the project, analysed the data, participated in the discussion of results, and wrote the manuscript. All the authors revised and approved the manuscript.

## Conflict of interest

The authors declare no conflict of interest.

## Funding

This work was supported by the PUE 040 grant (CONICET, Consejo Nacional de Investigaciones Científicas y Técnicas. Argentina), MINCYT (ex Ministerio de Ciencia y Tecnología) through the special grant “Ciencia y Tecnología contra el hambre” OC No RESOL-2021-289-APN-MCT/PROYECTO A12, ANPCyT (Agencia Nacional de Promoción Científica y Tecnológica. Argentina) PICT2019-0310, PICT2020-0362 to EW, and PICT2019 01916, PICT2020 0805 to RLC, Universidad Nacional del Litoral (CAI+D2020) and INTA 2023-PD-L01-I128 to SMS. JVC, RLC and EW are career members of CONICET; JC is a Fellow of the same Institution. SMS and JAMM are members of INTA; SMS is also a member of Universidad Nacional de Tucumán.

## Notes

### Competing Interest Statement

The authors have declared no competing interest.

